# Pan-cancer proteogenomic interrogation of the Ubiquitin Proteasome System

**DOI:** 10.64898/2026.03.23.713741

**Authors:** Tania J. González-Robles, Maha Khan, Paul Sastourné, Marisa Triola, Hua Zhou, Yuki Kito, Sharon Kaisari, David Fenyö, Gergely Rona, Yadira M. Soto-Feliciano, Benjamin G. Neel, Kelly V. Ruggles, Michele Pagano

## Abstract

Somatic mutations rewire the ubiquitin–proteasome system (UPS) to support tumor growth, but the proteome-wide consequences of cancer-driver alterations on UPS composition remain incompletely understood. Using harmonized proteogenomic data from up to 11 CPTAC cohorts, we performed an integrated pan-cancer analysis of UPS protein dysregulation, prognostic associations, and mutation-driven remodeling. We show that mRNA poorly predicts UPS protein abundance, that a defined set of E3 ligases is recurrently dysregulated across cancers, and that somatic mutations (most strikingly *TP53* loss) produce coherent UPS protein-quantitative trait locus (pQTL) signatures. Two case studies (UBR5 and TRIM28) illustrate orthogonal modes of UPS rewiring: a mutation-driven axis in which *TP53*-mutant tumors elevate UBR5 to support replication stress tolerance, and a lineage-driven axis in which TRIM28 engages tissue-restricted regulatory networks with opposing prognostic effects in glioblastoma versus head and neck cancer. Each axis exposes context-specific therapeutic vulnerabilities, including sensitivity to DNA damage response inhibitors (UBR5-high) and lineage-specific drug responses (TRIM28-high). Together, these analyses define a mechanistic framework for how cancer-driver mutations reshape proteostasis through the UPS and nominate mutation- and lineage-defined dependencies for precision degrader therapy. The harmonized pan-tissue atlas and the UbiDash interactive resource that underpin parts of this analysis are reported in our companion paper^1^.

**Highlights:**

- Proteome-centric profiling reveals UPS dysregulation invisible at the transcript level across human cancers
- Pan-cancer differential expression identifies recurrently dysregulated E3 ligases with prognostic value
- Genome-wide pQTL analysis uncovers mutation-driven UPS remodeling, with *TP53* loss as a dominant driver
- UBR5 and TRIM28 define orthogonal mutation- and lineage-driven axes of UPS rewiring with distinct therapeutic vulnerabilities

## Introduction

Cancer is fundamentally a disease of dysregulated proteins. Decades of genomic profiling have catalogued the somatic mutations that initiate and sustain malignancy^2–4^, yet the proteins encoded by mutated and wild-type alleles are the ultimate determinants of cellular phenotype, and their abundance is set as much by degradation as by synthesis. The ubiquitin–proteasome system (UPS) is one of the main routes of regulated intracellular proteolysis^5–8^, controlling the half-life of thousands of proteins involved in cell cycle progression, DNA-damage signaling, transcription, and metabolism^9–11^. Substrate selectivity in this system is conferred by E3 ubiquitin ligases (E3s) and reversed by deubiquitylases (DUBs), and the cooperative activity of these two enzyme classes defines which proteins are stabilized or eliminated in a given cellular context.

Somatic mutations are a hallmark of cancer^2–4^ and can affect the UPS by altering the activity of E3s or DUBs, thereby disrupting normal proteostasis^12,13^. Such perturbations can result in aberrant stabilization of oncogenic proteins or degradation of tumor suppressors. For instance, mutations in genes encoding E3s (e.g., *VHL*, *FBXW7*, *SPOP*, *KEAP1*) have been linked to altered UPS activity in specific tumor contexts. However, how recurrent cancer-driver mutations reshape UPS composition and function across tumor types, and how this influences E3-substrate relationships in a tissue- and lineage-specific manner, remains poorly understood. While individual examples of UPS dysregulation in cancer are well documented^9,11,12,14–18^, the proteome-wide effects of somatic mutations on UPS composition across tumor types remain largely unknown.

Despite this density of cancer-relevant biology, the proteome-level consequences of somatic mutations on UPS architecture remain largely uncharted. The human genome encodes over 600 E3s and substrate receptors and approximately 100 DUBs^19^. Nearly 20% of annotated cancer-driver genes encode UPS components^12,13^, yet many pan-cancer studies of UPS dysregulation have relied on transcript-level readouts that incompletely capture E3 abundance, complex assembly, and substrate engagement. Defining how recurrent driver mutations remodel the UPS at the protein level (and which E3-substrate relationships are rewired in a tumor- and lineage-specific manner) is therefore a pre-requisite for understanding proteostatic vulnerabilities in cancer, and for the rational use of the UPS as a therapeutic entry point, including emerging modalities describing targeted protein degradation^20–26^.

Here, we performed an integrated pan-cancer proteogenomic analysis of the UPS, focusing on E3s as regulators of substrate fate and potential therapeutic targets. By coupling differential protein expression, survival association, and genome-wide protein quantitative trait locus (pQTL) analyses across up to 11 CPTAC cohorts, we provide new insights into how somatic mutations rewire proteostasis through the UPS and identify two orthogonal axes: 1) a mutation-driven UBR5 axis active in *TP53*-deficient tumors and 2) a lineage-driven TRIM28 axis with opposing prognostic effects in glioblastoma and head and neck cancer; each exposing context-specific therapeutic vulnerabilities. The harmonized pan-tissue UPS atlas and the UbiDash interactive platform that underpin the resource layer of this analysis are described in our companion manuscript^1^.

## Results

### Divergence of RNA–protein establishes the need for proteome-centric UPS analysis

Although transcriptomic profiling remains widely used for functional inference, mRNA abundance is an imperfect surrogate of protein levels. To quantify the extent to which transcript levels reflect functional proteomic states in tumors, we analyzed the matched mRNA and protein abundance profiles across CPTAC cohorts (**Figure 1A,B**). For each sample and gene, we computed pairwise correlations between data modalities, enabling both sample-wise and gene-wise assessment of mRNA-protein expression concordance (**Supplementary Figure 1A,B**; see **Methods**). We observed a general positive association with a significant number of negative mRNA-protein expression correlations (**Figure 1C, Supplementary Table 1**), consistent with previous studies^27–29^. Gene set enrichment of genes with the lowest concordance between mRNA and protein (ρ < −0.2; *P* < 0.01) was the most discordant in regulatory pathways that operate downstream of transcription, including differential translation efficiency and protein half-lives, none of which could be inferred from mRNA alone (**Figure 1D,E and Supplementary Figure 1E**). Metabolic and housekeeping genes were overrepresented in those with the highest correlation (**Supplementary Figure 1C**), whereas transcription factors, signaling molecules, and UPS components frequently displayed weak or even inverse relationships (**Supplementary Figure 1D**). The high variability in mRNA-protein concordance among UPS components (**Figure 1F,G**) is consistent with proteins, rather than transcripts, serving as the direct biochemical effectors of UPS-mediated degradation (**Figure 1H**). This underscores the value of proteome-centered profiling to accurately characterize UPS biology and identify context-dependent vulnerabilities relevant to cancer progression and therapeutic targeting.

**Figure 1.**
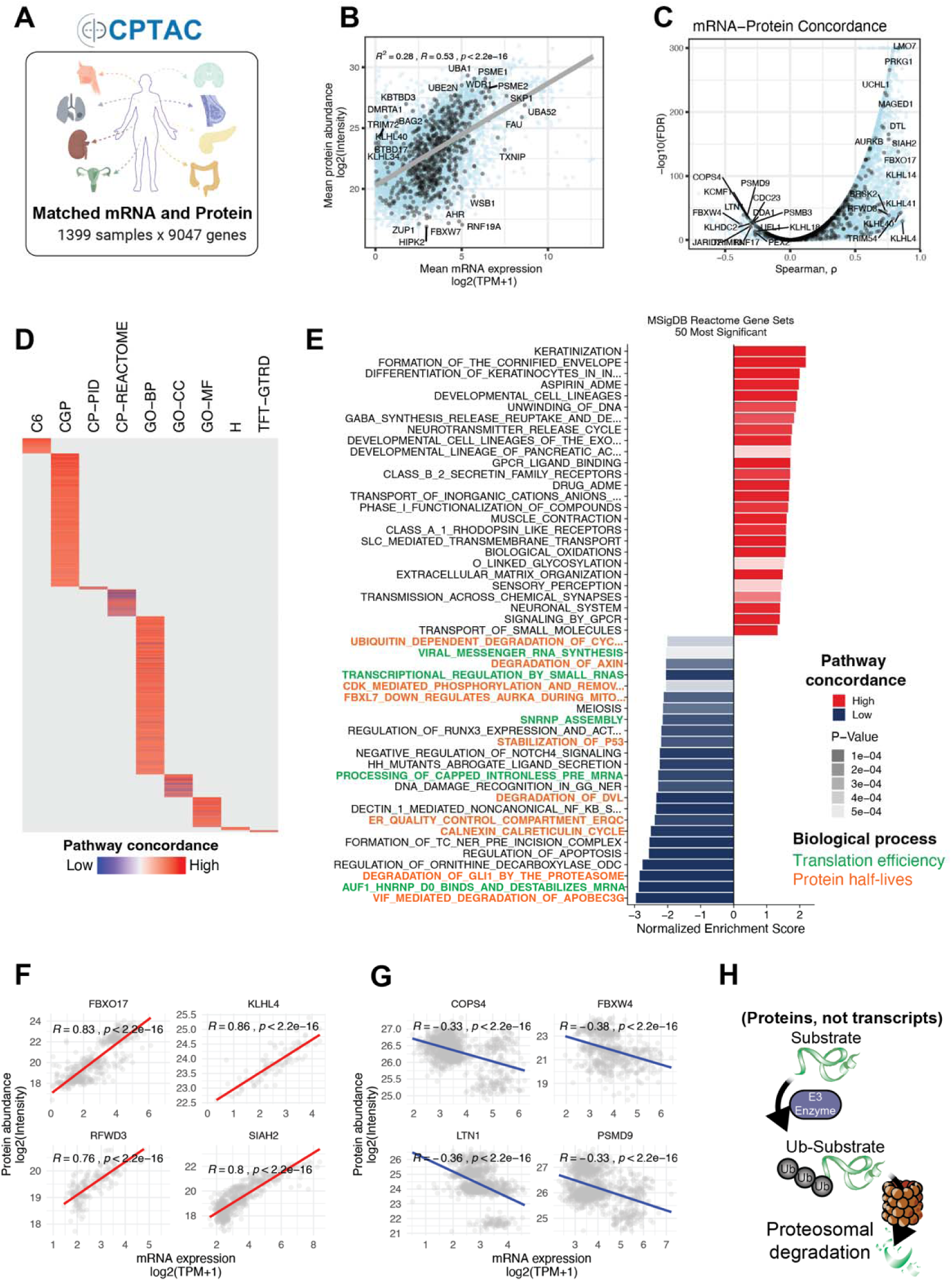
mRNA–protein divergence motivates a proteome-centric UPS analysis. (A) Overview of CPTAC cohorts and data modalities used for matched mRNA–protein correlation analysis across human tumors. (B) Scatter plot of mean mRNA expression versus mean protein abundance for genes quantified across CPTAC cohorts, highlighting UPS-associated genes in black. (C) Distribution of gene-wise mRNA–protein concordance measured by Spearman’s correlation; UPS-associated genes are highlighted in black. (D) Heatmap summarizing pathway-level mRNA–protein concordance based on gene-set Spearman’s correlations across multiple pathway databases (red, higher concordance; blue, lower concordance). (E) Bar plot showing the most significantly concordant (red) and discordant (blue) Reactome pathways (*P* < 0.05). (F) Representative UPS-associated genes with positive mRNA–protein Spearman’s correlations. (G) Representative UPS-associated genes with negative mRNA–protein Spearman’s correlations. (H) Schematic illustration of E3-mediated protein degradation, highlighting the central role of UPS proteins as direct effectors of proteome remodeling.

### Differential protein expression reveals UPS dysregulation in cancer

To characterize UPS dysregulation in cancer, we performed differential protein expression analysis across nine CPTAC cohorts, including eight matched tumor-normal adjacent tissues and one unmatched glioblastoma (GBM) dataset (**Figure 2A**). Among the 12,855 quantified proteins, 1,287 UPS components (comprising 670 E3s) were evaluated.

**Figure 2.**
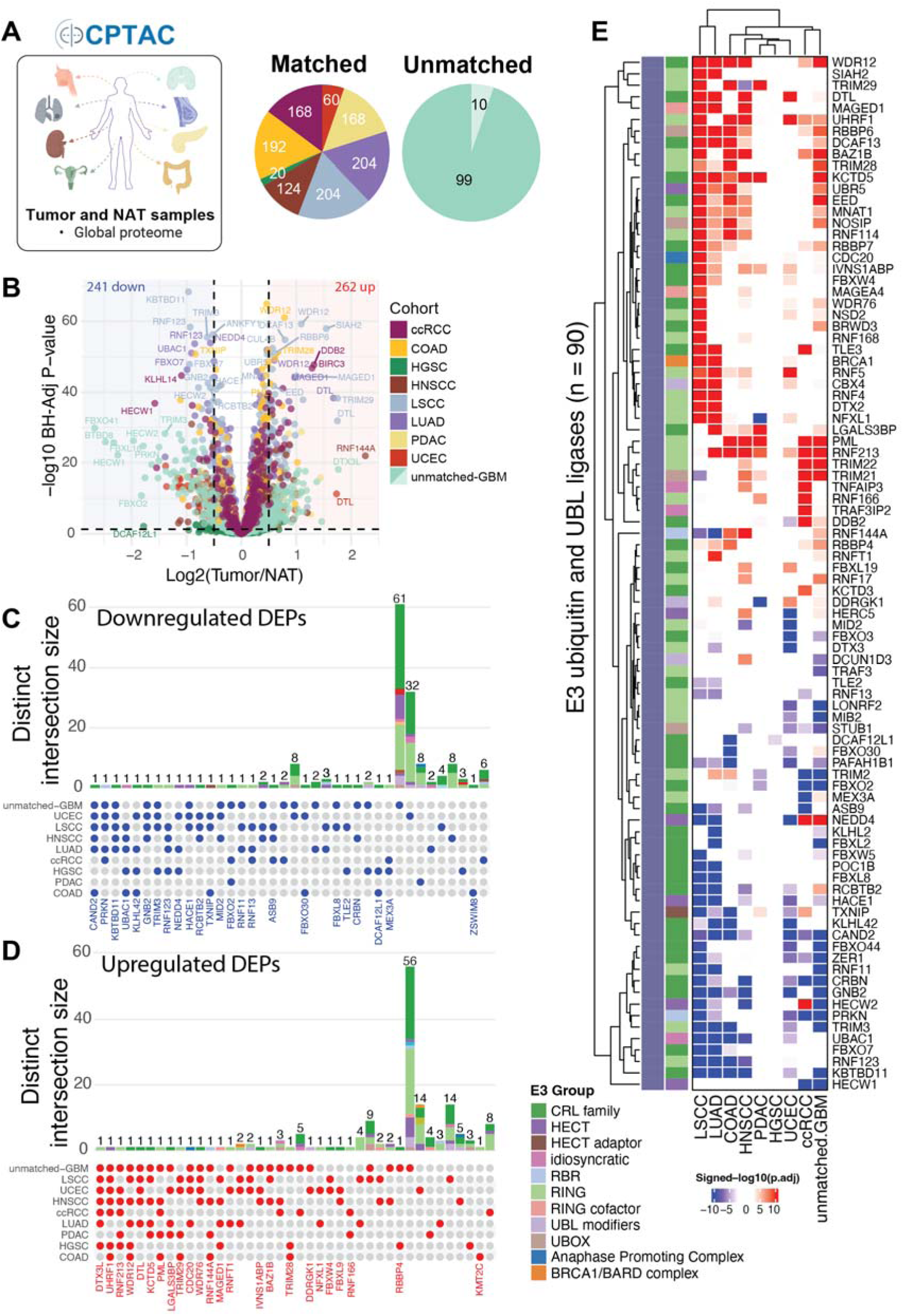
Tumor-specific dysregulation of UPS proteins across CPTAC cohorts. (A) Schematic illustrating CPTAC cohorts, sample composition, and proteomic data used for tumor-vs-normal differential expression (left), with the distribution of matched and unmatched normal tissues across cohorts (right). (B) Volcano plot summarizing differential protein abundance between tumor and normal tissues across CPTAC cohorts; horizontal dashed line, adjusted *P* < 0.05; vertical dashed lines, |log_2_FC| > 0.5. (C) UpSet plot summarizing UPS proteins significantly downregulated (adj. *P* < 0.05, |log_2_FC| > 0.5) in one or more cohorts. (D) UpSet plot summarizing UPS proteins significantly upregulated under the same thresholds. (E) Heatmap of selected UPS proteins (n = 90) showing significant up- (signed –log_10_(adj. *P*) > 2.5; red) or downregulation (< –2.5; blue) across CPTAC cohorts.

Across cohorts, 797 UPS-associated proteins were consistently quantified in at least 50% of CPTAC samples, including 410 E3s (**Supplementary Figure 2A,B**). Using a threshold of |log_2_ fold change (LFC)| > 0.5 and p-adj < 0.05, we identified 1,025 (562 unique) significantly dysregulated UPS proteins in at least one cohort, of which 503 (295 unique) were E3s (**Supplementary Figure 2C,D and Supplementary Table 2**). We evaluated the extent of UPS protein changes within individual cohorts and observed marked variation, ranging from 33 proteins in COAD (2.6%) to 357 in GBM (27.7%) (**Supplementary Figure 2C**). GBM, UCEC, and LSCC exhibited the greatest differential burden across the UPS and E3s, while COAD and HGSC showed minimal changes (**Figure 2B; Supplementary Figure 2C,D; Supplementary Table 2**). These differences were not due to missing values alone, as all cohorts were filtered to have similar protein coverage (797 UPS and 410 E3s quantified in at least 50% of samples) but reflect true variability in UPS protein changes across cancer types. Moreover, the elevated burden observed in GBM may be attributable to the absence of matched adjacent normal tissue, which can inflate differential expression estimates. This reflects an inherent limitation of sample availability, as the collection of normal adjacent brain tissue from patients with GBM is not ethically feasible.

Although UPS components were less frequently changed in abundance than the global proteome (Fisher’s exact test, odds ratio < 1, p-adj < 0.0001, **Supplementary Figure 2G**), consistent with the selective constraint on the core proteostasis machinery, a distinct subset showed significant and context-specific changes (**Figure 2B and Supplementary Figure 2A-D**). These patterns suggest that UPS dysregulation is selective rather than random. Specifically, several E3s displayed consistent dysregulation across four or more cancer types (**Figure 2C-E; Supplementary Figure 2E,F**, p-adj < 0.05, |LFC| > 0.5). These included downregulation of the tumor suppressor PRKN and putative tumor suppressor RNF123 (**Figure 2C,E; Supplementary Table 1**) and upregulation of oncogenic or pro-proliferative E3s, such as SKP2 and CDT2 (*DTL*) (**Figure 2D,E; Supplementary Table 1**). Notably, the protein levels of several canonical tumor suppressor E3s such as VHL and FBXW7 were frequently undetected, a pattern consistent with mutation-driven protein instability^30,31^, epigenetic silencing^32^, lack of tryptic peptides needed for MS detection^33^, or copy number loss^34^ (**Supplementary Figure 2H**).

This analysis also identified context-dependent changes. KEAP1, a Cullin-RING Ligase 3 (CRL3) substrate receptor and tumor suppressor in lung adenocarcinoma (LUAD), was downregulated in LUAD (LFC = −0.2, p-adj < 0.001), consistent with previous reports^35–38^, but was upregulated in LSCC (LFC = 0.4, p-adj < 0.001) and GBM (LFC = 0.5, p-adj < 0.001) (**Supplementary Table 2**). In LSCC, KEAP1 protein abundance did not correlate with predicted activity, as LSCC harbors frequent *KEAP1* mutations that cluster within the Kelch/DGR domain required for NRF2 substrate recruitment (**Supplementary Figure 2H**), likely rendering the elevated protein levels functionally inert^38^. To confirm KEAP1 loss of activity, we evaluated the anti-correlation between KEAP1 and its known target NRF2 (*NFE2L2*), as well as NRF2 downstream effectors (NQO1 and GCLC), but these proteins were not consistently quantified across CPTAC datasets. Differential expression also highlighted less-characterized UPS components with conserved suppression across tumors, such as KBTBD11 and CAND2, suggesting possible tumor suppressive roles. Conversely, NEDD4 and RNF144A showed heterogeneous tumor-type-specific patterns (**Figure 2E**).

Overall, our differential UPS protein analysis revealed both pan-cancer- and tissue-specific dysregulation. Given the catalytic nature of E3s, it is important to note that even modest shifts in their abundance can produce quantitative effects on substrate stability and downstream signaling, underscoring the need for protein-level profiling to accurately identify UPS vulnerabilities in cancer.

### UPS protein expression levels stratify cancer prognosis in a tissue-specific manner

To assess the clinical consequences of UPS dysregulation, we stratified patients by UPS-associated protein abundance and examined their relationship with patient survival across 11 CPTAC tumor types (**Figure 3A**). For each of the 922 UPS proteins (including 411 E3s), we classified patients within each tumor type into UPS-high or UPS-low groups, based on the upper and lower quartiles of normalized protein abundance (**Supplementary Figure 3A,B**). Stratification was performed using both pan-cancer and per-cancer approaches, followed by systematic survival analysis comparing overall survival between the UPS-high and UPS-low groups (**Figure 3B,C**).

**Figure 3.**
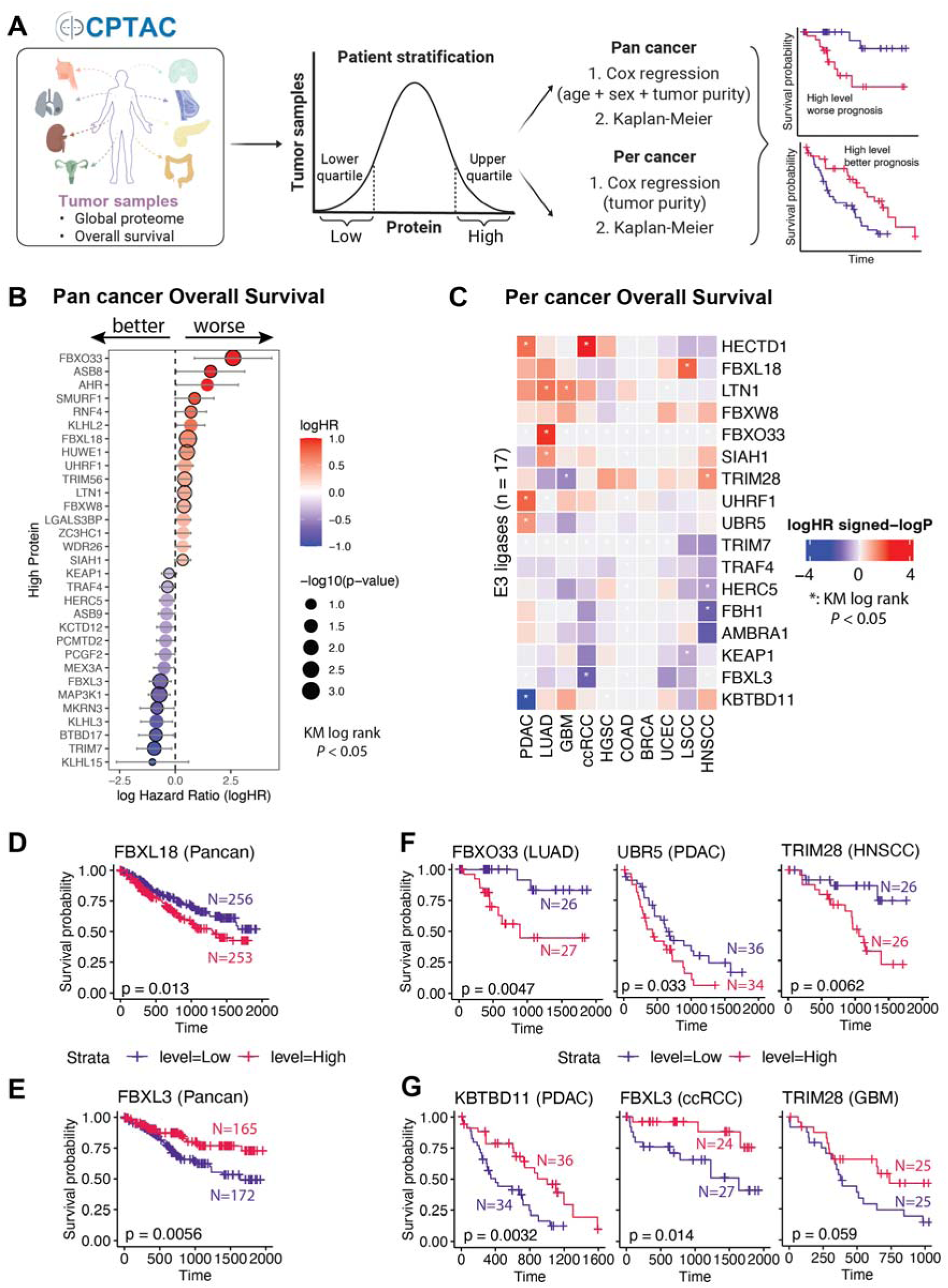
Overall-survival profiling of UPS-associated proteins. (A) Schematic of the survival analysis design (UPS-high vs UPS-low quartile stratification, KM and Cox models). (B) Bubble plot of pan-cancer Cox log-hazard ratios for UPS-associated proteins; red, higher abundance associated with worse survival; blue, better; size, statistical significance. (C) Heatmap of per-cancer Cox log-hazard ratios across tumor types; asterisks, significant after multiple-testing correction. (D) Kaplan–Meier curve for a representative UPS protein with poorer pan-cancer survival when high. (E) KM curve for a representative protein with better pan-cancer survival when high. (F,G) KM curves of representative lineage-specific UPS-associated proteins linked to worse (F) or better (G) survival in selected tumor types.

Kaplan–Meier survival analysis identified 153 E3s whose protein expression was significantly associated with overall patient survival in at least one cohort (log-rank *p* < 0.05, **Supplementary Figure 3C**). Multivariate Cox regression models, adjusting for age, sex, and tumor purity (pan-cancer) or for tumor purity alone (per-cancer), confirmed independent prognostic association of multiple E3s (|logHR| > 0.5, *p* < 0.05, log-rank *p* < 0.05; **Figure 3B,C; Supplementary Figure 3D,E; Supplementary Table 1**). This dual-test framework, with effect-size filtering, prioritizes robust associations while mitigating false positives.

Pan-cancer analysis identified relatively few consistent E3 survival predictors. Notably, high expression of FBXL18 was significantly associated with worse patient survival, likely driven by effects in LSCC (**Figure 3D; Supplementary Figure 3D**), whereas an increased abundance of FBXL3 predicted improved clinical outcomes (**Figure 3E**). Both proteins are members of the F-box protein family and function as substrate receptor adaptors within the Cullin-RING Ligase 1 (CRL1) E3 complexes^19^. In contrast, a larger number of E3s displayed tumor-specific prognostic effects. High expression of FBXO33 and UBR5 predicted poor prognosis in LUAD and PDAC, whereas overexpression of KBTBD11 and FBXL3 was associated with improved survival in PDAC and ccRCC, respectively (log-rank *p* < 0.05, **Figure 3F,G**). Interestingly, high TRIM28 levels predicted improved survival in LUAD (*p* = 0.031) and a trend (*p* = 0.059) toward a favorable outcome in GBM, yet showed poor survival in HNSCC (**Figure 3F,G; Supplementary Figure 3E**).

Together, these findings reveal that E3s can serve as both prognostic biomarkers and mechanistic indicators of tumor-specific UPS dependency.

## Cancer-associated mutations reshape the UPS protein landscape

To determine whether cancer-associated mutations alter UPS protein abundance, we modeled each UPS protein (Y) as a function of mutation status (M) while controlling for tumor purity (P) and patient cohort (C) (**Supplementary Figure 4A and Methods, Eq. 1**). For this purpose, we analyzed 9,495 recurrent gene mutations that were detected in at least 11 tumor samples across 10 CPTAC cohorts to avoid unbalanced or underpowered comparisons (**Supplementary Figure 4B**).

Systematic UPS protein quantitative trait locus (pQTL) analysis revealed 17,925 significant UPS protein–gene mutation associations (false discovery rate or FDR < 10%; **Figure 4A; Supplementary Table 1**), demonstrating widespread mutation-associated proteomic restructuring across all UPS levels, including E2s (e.g., UBE2T), E3s (e.g., DDB2, CDC20, FBXO22, UBR5, CDT2 (*DTL*), TRIM3, and TRIM29), deubiquitylases (e.g., USP28 and MINDY1), and proteasome-associated proteins (e.g., AURKB and PLK1) (**Figure 4A,B and Supplementary Table 1**). These alterations have been observed across multiple cancer types and likely reflect heterogeneous regulatory mechanisms. For example, TRIM29 showed coordinated upregulation at both mRNA and protein levels (**Supplementary Figure 4C-F; Supplementary Figure 5**), whereas UBR5 exhibited a protein-specific expression phenotype consistent with post-transcriptional and/or post-translational regulation (**Supplementary Figure 4C-F, Supplementary Figure 6**)^39^. *TP53* mutations produced a striking proteomic signature (**Figure 4A,B; Supplementary Table 1**). Tumors with *TP53* point mutations showed significant dysregulation in 111 proteins (FDR < 0.05) with upregulation in CDT2 (*DTL*), CDC20, and PLK1 (components promoting S-phase progression and mitotic bypass) and downregulation of TP53 transcriptional targets, such as DDB2 and FBXO22, consistent with impaired DNA repair signaling (**Figure 4B,C**). These observations align with known TP53 biology and reveal broader effects on UPS regulation.

**Figure 4.**
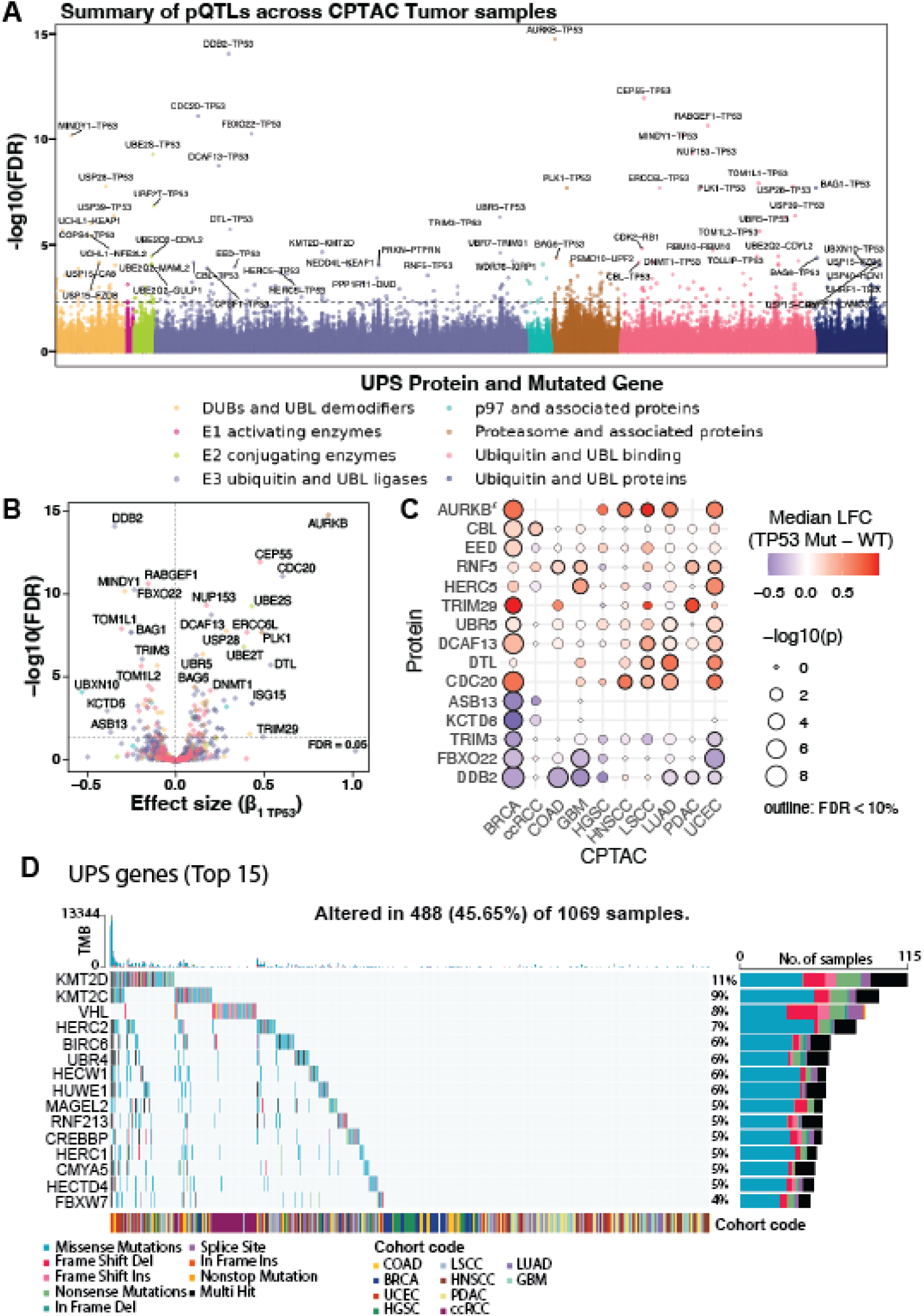
Mutation-to-protein quantitative-trait-locus (pQTL) analysis of UPS remodeling. (A) Manhattan plot of significant somatic-mutation–UPS-protein associations across CPTAC cohorts; y-axis, transformed FDR; horizontal dashed line, FDR = 0.001. (B) Volcano plot of *TP53* mutation effects on UPS protein abundance; x-axis, β₁; horizontal dashed line, FDR = 0.05. (C) Bubble plot of median log_2_ fold changes of selected UPS proteins in *TP53*-mutant vs wild-type tumors across cohorts; bubble size, Wilcoxon *P*; black outline, FDR < 10%. (D) Oncoplot of the 15 most frequently mutated UPS-associated genes across CPTAC cohorts with tumor mutational burden (TMB).

Several E3s exhibited mutation-dependent and context-specific regulation. Both mRNA and protein expression of TRIM29 was consistently elevated in *TP53*-mutant BRCA and PDAC (Wilcoxon *P* < 0.05; **Figure 4C; Supplementary Figure 5**), suggesting that observed changes are driven by transcriptional regulation associated with *TP53* mutations. UBR5 protein, a HECT-type E3 involved in DNA damage response and chromatin maintenance^40–43^, was consistently upregulated in *TP53*-mutant tumors across seven cohorts (BRCA, COAD, UCEC, HGSC, HNSCC, LSCC, and LUAD; Wilcoxon *P* < 0.05, **Figure 4C, Supplementary Figure 6B-D**). Of these, UBR5 upregulation observed in 3 out of 7 cohorts (COAD, LSCC and LUAD) was likely driven by transcriptional activity (**Supplementary Figure 6A**), while 4 out of 7 tumors (BRCA, UCEC, HGSC and HNSCC) only showed post-translational changes (**Supplementary Figure 6B**). UBR5 is known to preferentially assemble K48-linked polyubiquitin chains, which directly signal proteasomal degradation^40^, thus providing a mechanistic rationale linking UBR5 elevation to enhanced substrate turnover in *TP53*-deficient contexts. This finding is consistent with the proposed UBR5 oncogenic function in *TP53*-deficient cancers^44^. In contrast, FBXO22, an E3 protein transcriptionally induced by TP53^45^, was downregulated in *TP53*-mutant tumors (**Figure 4B,C**), reflecting a loss of TP53-mediated transcription.

UPS genes also harbored frequent somatic mutations (**Figure 4A,D; Supplementary Table 1**). Canonical tumor suppressors, such as *VHL* (8%) and *FBXW7* (4%), exhibited high mutation frequency, consistent with their well-established roles in substrate stabilization and tumorigenesis upon loss^30,31,46^. Less-characterized E3s, including *RNF213* (5%), were also found to be recurrently mutated across cancer types, suggesting broader roles in proteostasis rewiring. RNF213 has been implicated in the regulation of hypoxia-induced inflammatory cell death in cancer^47^ and is broadly amplified across tumors in cBioPortal (www.cbioportal.org, data not shown). This was consistent with the elevated RNF213 protein abundance observed earlier in CPTAC samples (**Figure 2E**). These results demonstrate that somatic mutations alter distinct UPS remodeling axes in cancer, revealing mutation-driven proteostatic states with potential therapeutic relevance.

### Distinct UPS remodeling axes exemplified by UBR5 and TRIM28

To understand the functional consequences of UPS remodeling, we investigated two representative E3s (UBR5 and TRIM28) that exhibited strong but divergent cancer associations. We evaluated how variation in UPS protein abundance is related to mutational context and tumor lineage. Associated functional programs were also characterized using four complementary approaches: 1) UPS protein co-regulation analysis, 2) proteome-wide pathway enrichment, 3) lineage-specific dependency profiling^48,49^, and 4) drug sensitivity analysis^50^ (**Figure 5A, Supplementary Figure 7 and Supplementary Table 3**).

**Figure 5.**
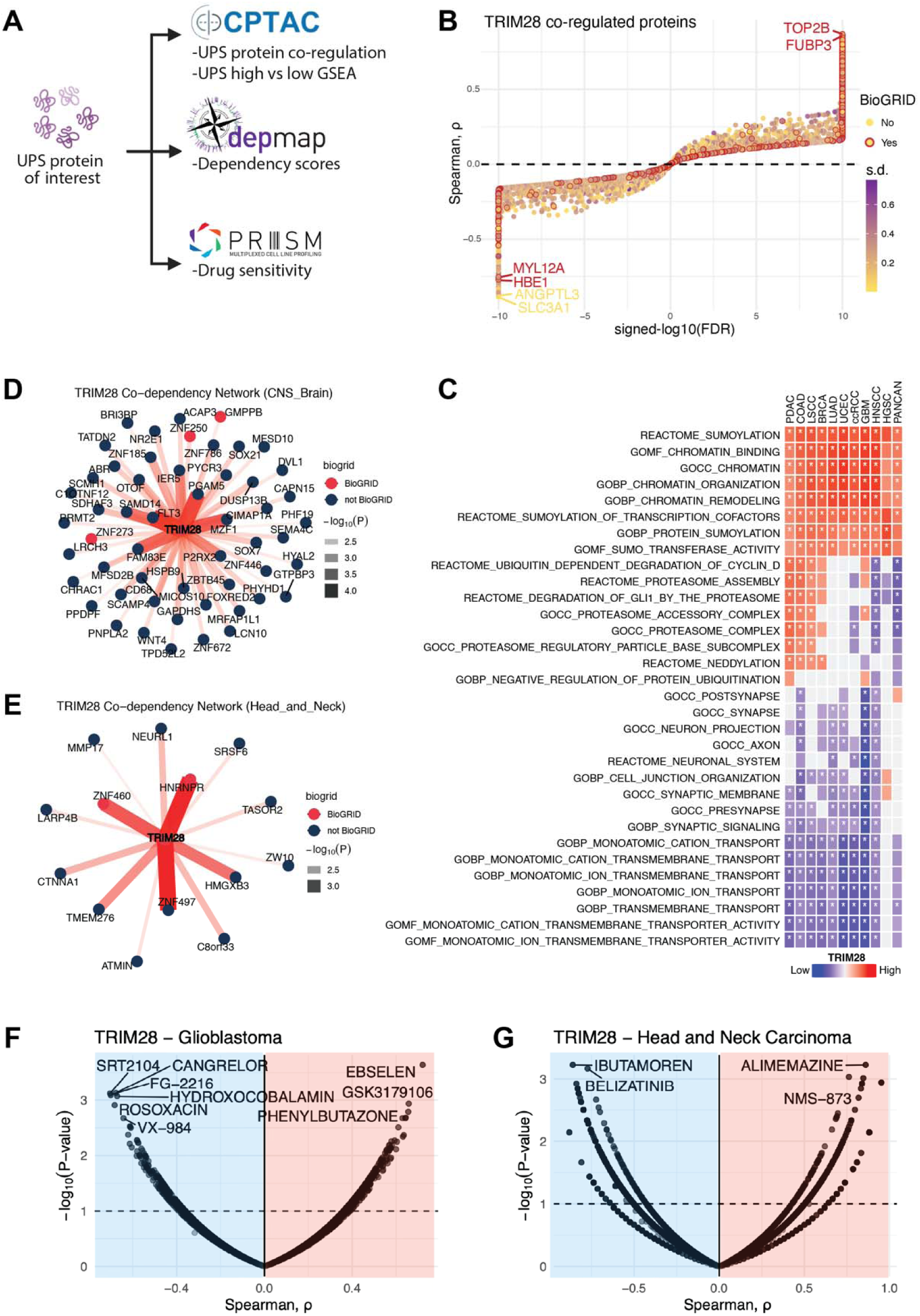
TRIM28 exemplifies lineage-associated UPS vulnerabilities. (A) Schematic of the integrated analysis: protein co-regulation, pathway enrichment, co-dependency, and drug sensitivity for UBR5 and TRIM28. (B) TRIM28 protein co-regulation pan-cancer; color, between-cohort SD of correlation; red outline, BioGRID interactors; axes, signed transformed FDR and ρ. (C) GSEA of TRIM28 co-regulated proteins (preranked, MSigDB, 1,000 permutations). (D) TRIM28 co-dependency network in CNS/brain cell lines (DepMap CRISPR Chronos); nodes, top positively co-dependent genes; red, BioGRID interactors. (E) TRIM28 co-dependency network in HNSCC cell lines, plotted as in (D). (F) TRIM28-protein–PRISM compound associations in glioblastoma cell lines; horizontal dashed line, FDR = 10%. (G) TRIM28-protein–PRISM compound associations in HNSCC cell lines.

#### UBR5 defines a mutation-associated UPS remodeling axis

UBR5 protein abundance was consistently elevated in *TP53*-mutant tumors across multiple cancer types (**Supplementary Figure 6B**) and high UBR5 expression was associated with poor survival in PDAC (**Figure 3F**). Notably, UBR5 protein upregulation was not consistently accompanied by a proportional increase in mRNA expression, resulting in significantly increased protein-to-mRNA ratios across cancers (Wilcoxon *P* < 0.05, **Supplementary Figure 8**). Therefore, we interpreted this pattern as post-transcriptional regulation, consistent with increased UBR5 protein stability or reduced turnover, rather than transcriptional induction.

Pathway enrichment analysis of UBR5-high tumors revealed consistent upregulation of DNA replication and repair machinery, checkpoint regulation, chromatin remodeling factors, downregulation of TP53 target genes, and AKT-driven proliferative signaling pathways (**Supplementary Figure 9B,C and Supplementary Table 3**). These programs align with the known biochemical roles of UBR5 in genome maintenance as well as cell cycle regulation and suggest that elevated UBR5 supports tolerance to replication stress in *TP53*-deficient contexts^42,51,52^.

To assess the functional dependencies associated with this state, we performed DepMap co-dependency analysis (**Supplementary Figure 7F, 9D, 10A; Supplementary Table 3**). The UBR5-high cell lines exhibited coordinated dependency patterns with genes involved in replication-coupled chromatin maintenance and DNA damage response pathways, including TRIP12 and MED25, along with chromatin- and ubiquitin-linked stress regulators (**Supplementary Figure 9D, 10A and Supplementary Table 3**).

Consistent with these dependencies, PRISM (profiling relative inhibition simultaneously in mixtures) drug sensitivity profiling showed that UBR5-high cell lines were more sensitive to compounds targeting RNA splicing, cell cycle regulation and PI3K/mTOR signaling (PRMT5 inhibitor JNJ-64619178), as well as innate immune signaling (STING agonist MIW-815) (**Supplementary Figure 9E; Supplementary Table 3**). Together, these findings suggest that UBR5 is a mutation-associated regulator of proteostasis, whose elevation supports genome maintenance programs while simultaneously creating exploitable therapeutic vulnerabilities.

#### TRIM28 defines a lineage-specific UPS remodeling axis

In contrast to UBR5, TRIM28 abundance was not associated with recurrent somatic mutations but instead displayed lineage-specific phenotypes. Protein co-regulation analysis revealed that TRIM28 abundance was positively associated with chromatin- and transcription-linked factors (e.g., TOP2B and FUBP3) and negatively associated with cytoskeletal and metabolic proteins (**Figure 5B**). This pattern suggests that TRIM28 participates in coordinated regulatory programs with its positively associated partners and may contribute to the selective suppression of negatively associated proteins. Pan-cancer analyses revealed that TRIM28-high tumors were enriched in MYC- and E2F-driven transcriptional programs, RNA processing factors, and chromatin regulatory complexes (**Figure 5C; Supplementary Table 3**). However, the clinical consequences of TRIM28 upregulation diverged sharply by tissue context, with high expression correlating with a trend to improved survival in GBM but poor survival in HNSCC (**Figure 3F,G**).

To understand these opposing associations, we performed context-specific pathway enrichment analysis. In GBM, TRIM28-high tumors showed increased expression of DNA repair and chromatin organization pathways, consistent with a genome-stabilizing role (**Figure 5C; Supplementary Table 3**). In contrast, TRIM28-high HNSCC tumors were enriched in mitochondrial metabolism and immune mimicry programs, whereas cell adhesion and EMT programs appeared to be downregulated (**Figure 5C; Supplementary Table 3**), suggesting distinct functional roles across lineages.

The lineage-resolved dependency analyses reinforced these distinctions. In CNS-derived cell lines, including GBM cell models, TRIM28 dependency was correlated with growth and stress adaptation programs, including RTK-PI3K signaling (FLT3), mitochondrial stress regulation (PGAM5), transcriptional reprogramming (MZF1), and EGFR-MAPK pathway scaffolding (FAM83E) (**Figure 5D; Supplementary Figure 10B; Supplementary Table 3**). In contrast, head and neck cancer cell lines exhibited TRIM28-associated dependencies enriched for chromatin and RNA regulatory machinery, including KRAB-zinc finger transcription factors (ZNF460 and ZNF497) and the RNA-binding protein HNRNPR, consistent with lineage-specific transcriptional repression and RNA processing programs (**Figure 5E; Supplementary Figure 10B; Supplementary Table 3**).

Drug sensitivity profiling^50^ further reinforced the lineage-specific nature of TRIM28 dependence. In the CNS, TRIM28-high cell lines were preferentially sensitive to compounds targeting chromatin-associated DNA damage regulation and stress adaptation, including the SIRT1 activator SRT2104 and DNA damage response signaling through the DNA-PK inhibitor VX-984^53^, while exhibiting relative resistance to redox and inflammatory agents, such as ebselen and phenylbutazone, respectively (**Figure 5F; Supplementary Table 3**). In contrast, TRIM28-high HNSCC cell lines displayed increased sensitivity to drugs linked to metabolic and growth signaling, including ibutamoren and belizatinib, while showing resistance to compounds associated with proteostasis, cytoskeletal remodeling, and EMT-related programs, such as the VCP/p97 inhibitor NMS-873 and GPCR modulator alimemazine (**Figure 5G; Supplementary Table 3**). These results indicate that TRIM28 reshapes the local proteostatic and signaling environment in a strong lineage-dependent manner.

Together, UBR5 and TRIM28 define two orthogonal modes of UPS remodeling: 1) a mutation-driven axis, in which *TP53* mutations allow for increased UBR5 abundance to support replication stress tolerance and chromatin maintenance, and 2) a lineage-driven axis, in which TRIM28 engages distinct regulatory networks to shape tumor behavior in a tissue-specific manner (**Figure 6**). These complementary mechanisms illustrate how selective modulation of UPS components can support cancer progression while simultaneously exposing context-specific therapeutic opportunities.

**Figure 6.**
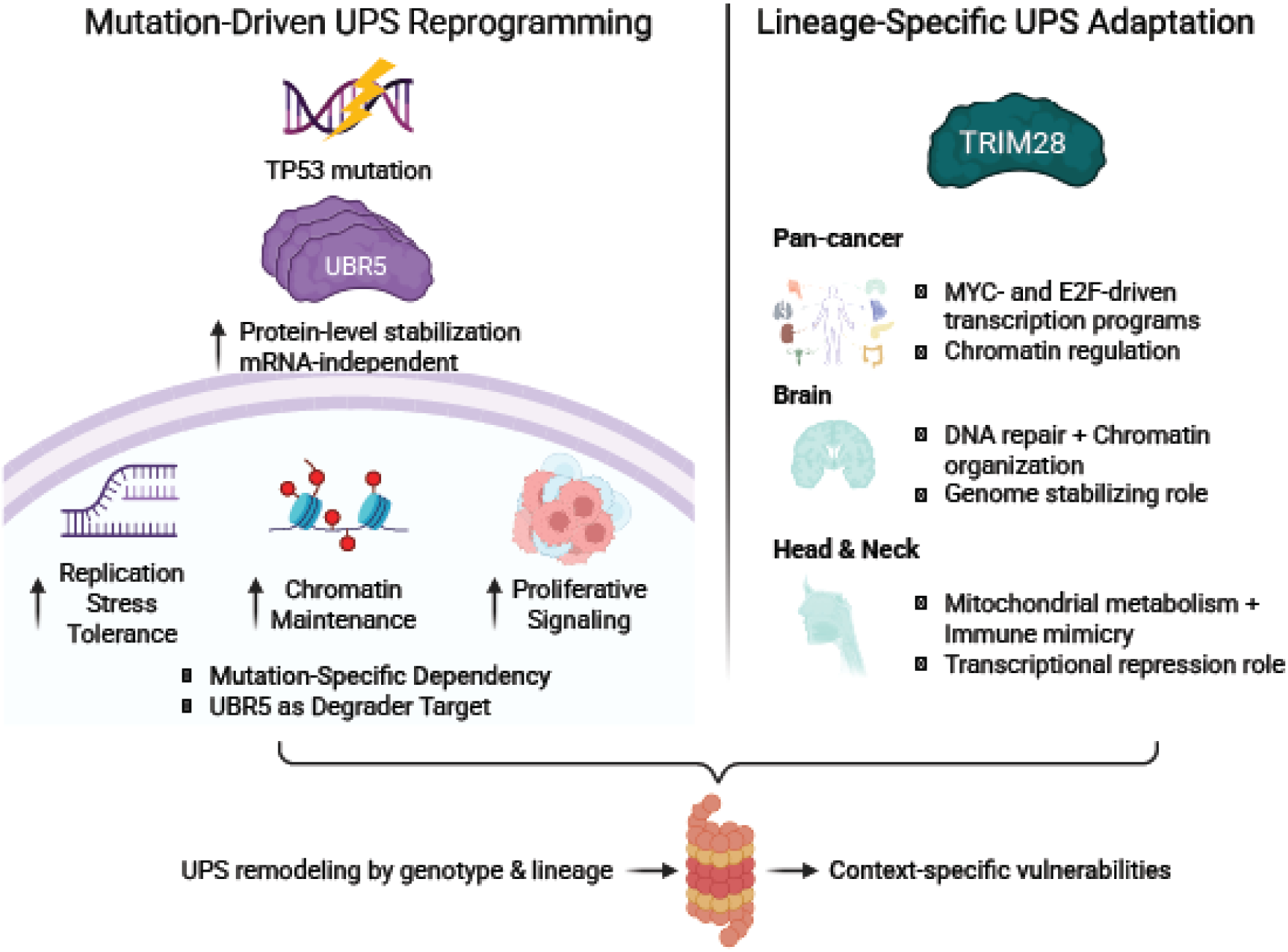
Conceptual model of mutation- and lineage-driven UPS remodeling in cancer. Schematic illustrating two orthogonal modes of UPS regulation in tumors: a mutation-driven axis, exemplified by *TP53*-driven UBR5 elevation supporting replication stress tolerance and chromatin maintenance (left), and a lineage-driven axis, exemplified by TRIM28 engaging tissue-restricted regulatory networks to reshape proteostasis and signaling (right), generating distinct therapeutic vulnerabilities (DDR/PI3K inhibitor sensitivity in UBR5-high *TP53*-mutant tumors; lineage-specific drug responses in TRIM28-high CNS vs HNSCC).

## Discussion

Mutation- and lineage-driven remodeling of the UPS is a pervasive but underappreciated dimension of cancer proteostasis. By integrating differential protein expression, genome-wide pQTL mapping, and survival association across up to 11 CPTAC cohorts, this study defines (at the proteome level) how somatic alterations reshape the abundance and stoichiometry of UPS components, including E3s and DUBs in human tumors. Two orthogonal axes emerge from this analysis: 1) a mutation-driven axis, exemplified by UBR5 activation in TP53-mutant tumors, and 2) a lineage-driven axis, showing opposing prognostic effects of TRIM28 across tissues.

Together, these data illustrate that UPS dysregulation in cancer is neither uniform nor purely transcriptional, and that proteogenomic resolution is essential to capture it. The harmonized pan-tissue atlas and the UbiDash interactive resource that complement these mechanistic analyses are described in our companion manuscript^1^.

Each tissue maintains a characteristic UPS landscape shaped by physiological demands, and tumors frequently co-opt for or distort these circuits. Squamous cancers, for instance, elevate stress-handling E3s^54,55^, while glandular tumors upregulate replication-coupled CRLs^56–58^, and tumor-suppressive ligases, such as FBXW7, are commonly mutated in various cancers^30,31^.

Although many changes are moderate in magnitude, the catalytic nature of E3 ligases means that small shifts can have large effects on substrate turnover and signaling. These findings underscore the utility of proteomic profiling for revealing UPS remodeling, which is not always detectable at the transcript level. Recurrent dysregulation of DUBs, such as USP28, further underscores an additional regulatory layer that warrants deeper, dedicated investigation in future studies.

Somatic mutations further reshape the UPS by integrating transcriptional, post-transcriptional, and post-translational regulation. *TP53*-mutant tumors exemplify this principle, displaying coordinated upregulation of UPS-associated proliferative drivers (CDT2 and CDC20) and downregulation of genome-stabilizing E3s (DDB2 and FBXO22). In agreement with prior studies, UBR5 has emerged as a prominent mutation-responsive E3, recurrently upregulated in *TP53*-mutant tumors and associated with poor prognosis^59–62^. Its elevation at the protein, rather than mRNA, level suggests post-translational stabilization and highlights a potential synthetic-lethal or degrader target in *TP53*-deficient cancers.

In contrast, TRIM28 showed lineage-driven UPS remodeling. Although not mutation-associated, TRIM28 exhibits divergent functional roles across tissues: in brain cancer, it correlates with chromatin repression and improved outcomes, whereas in head and neck cancer, it aligns with enhanced translation, metabolic activity, and poor survival. These differences are reinforced by lineage-specific dependency and drug-sensitivity profiles, emphasizing the need to evaluate UPS components within their tissue context.

Taken together, the analyses reported here demonstrate the UPS as a structured, mutation-responsive, and lineage-dependent layer of cancer proteostasis rather than a static catalogue of housekeeping enzymes. The UBR5 and TRIM28 axes provide concrete examples of how genotype and tissue identity converge on individual ligases to produce distinct dependencies and prognostic outcomes (**Figure 6**), and the genome-wide pQTL map presented here position many additional E3-substrate relationships for hypothesis-driven follow-up. Orthogonal experimental validation of mutation-conditional E3-substrate edges, together with single-cell and spatial proteomics in matched cohorts, will be required to convert these proteogenomic associations into mechanistic and therapeutic gains. The harmonized pan-tissue atlas and UbiDash interface described in our companion paper^1^ make the underlying data queryable and extendable by the community.

The UPS underpins most targeted protein degradation strategies; however, how cancer-driver mutations remodel UPS composition at the proteome level remains incompletely understood. By integrating proteogenomic profiles across multiple CPTAC cohorts, we provide a comprehensive map of UPS protein dysregulation, mutation-associated rewiring, and lineage-specific dependencies in human cancers. We show that E3 abundance is highly context-dependent, that oncogenic mutations such as *TP53* loss reshape degradative circuits at the protein level, and that ligases such as UBR5 and TRIM28 define distinct axes of mutation-driven and lineage-driven UPS remodeling. This study identifies tumor-selective ligases suitable for therapeutic hijacking, exposes emergent proteostasis vulnerabilities, and provides a mechanistic foundation for the design of next-generation degraders, complementing the harmonized pan-tissue atlas and UbiDash resource described in our companion paper^1^.

This study provides a comprehensive view of mutation- and lineage-driven UPS remodeling in cancer. However, several limitations should be acknowledged. Our analyses rely on bulk proteogenomic data, which lack single-cell or spatial resolution and therefore cannot capture microenvironmental heterogeneity or cell-type-specific UPS remodeling. The CPTAC datasets analyzed also overrepresent female-specific tumor types (e.g., breast, endometrial, and ovarian) while underrepresenting male-specific cancers, such as prostate adenocarcinoma, which may limit generalizability. In addition, the technical constraints of proteomic profiling, particularly missing values and bias toward highly abundant proteins, may hinder the detection of low-abundance or tissue-restricted ligases. Moreover, many E3s regulate their own stability through auto-ubiquitylation, such that observed protein-level changes may partially reflect altered self-turnover dynamics rather than solely transcriptional or translational regulation, an effect that could also contribute to the observed mRNA-protein discordance. Finally, survival associations were based on overall survival data without accounting for treatment history or disease stage, and bulk profiling could not resolve temporal or therapy-induced dynamics. Future studies incorporating longitudinal, single-cell, and perturbation-based datasets will be essential to refine these insights and to validate predicted degradative interactions for translational applications in targeted protein degradation.

## Supporting information

Key Resources

## Resource Availability

See Key Resources Table.

## Lead contact

Further information and requests for resources should be directed to and will be fulfilled by the lead contacts, Kelly V. Ruggles and Michele Pagano.

## Materials availability

This study did not generate new unique reagents.

## Data and Code Availability Statement

The gene list annotating UPS components was downloaded from https://www.proteostasisconsortium.com/pn-annotation/ on June 22, 2024. Raw and processed proteomics data, as well as open-access genomic data, can be obtained via Proteomic Data Commons (PDC) at https://pdc.cancer.gov/pdc/cptac-pancancer. Raw genomic and transcriptomic data files can be accessed via the Genomic Data Commons (GDC) Data Portal at https://portal.gdc.cancer.gov with dbGaP Study Accession phs001287.v16.p6. Complete CPTAC pan-cancer controlled and processed data can be accessed via the Cancer Data Service (CDS: https://dataservice.datacommons.cancer.gov/). Additional data reported here can be shared by the lead contacts upon request. All original code was deposited at GitHub (https://github.com/ruggleslab/ubiDash) and is available interactively as the UbiDash R Shiny app (https://ruggleslab.shinyapps.io/UbiDash/), described in our companion manuscript^1^.

## Experimental Models and Subject Details

### Human subjects and clinical specimens

This study exclusively used de-identified human tumor and normal tissue proteogenomic datasets that were previously generated, peer-reviewed, and released by large consortia and public repositories. No new human subjects were recruited and no prospectively collected biospecimens were obtained by the authors for this work. Clinical annotation, sample procurement, and informed consent procedures for all CPTAC cohorts were conducted by the original contributing centers in accordance with the Declaration of Helsinki and local institutional review boards, and only data that has passed the corresponding data access and compliance checks were used. All CPTAC pan-cancer datasets were accessed through the NCI’s Proteomic Data Commons, Genomic Data Commons, and Cancer Data Service under approved dbGaP data use agreements (phs001287.v16.p6).

### Cancer cell lines

Cancer cell line proteomic and transcriptomic profiles, dependency (DepMap CRISPR), and drug-response (PRISM) data were obtained from previously published peer-reviewed resources. No new cell lines were generated or experimentally manipulated in this study.

### Inclusion and exclusion criteria

For corresponding analyses, we restricted samples to those with matched proteomic and genomic data (e.g., CPTAC pan-cancer cohorts with both mutation and protein profiles available) and excluded samples failing consortium-level quality control or lacking key clinical annotations.

## Method Details

### Differential protein expression analysis

Protein-level differential expression was performed using DEqMS^63^, which builds on the Limma framework^64^. DEqMS improves statistical power by modeling the relationship between protein variance and the number of peptide-spectrum matches (PSMs); the spectraCounteBayes function estimates prior variance based on PSM counts and integrates this into variance moderation. All p-values were adjusted using the Benjamini–Hochberg procedure to control the false discovery rate.

#### Tumor vs. normal-adjacent tissue comparisons

Differential expression was assessed between tumors and matched normal-adjacent tissue samples across CPTAC cohorts. Proteins were filtered to retain those with sufficient coverage across paired samples. DEqMS was applied using sample-specific PSM counts and contrasts were independently computed for each cohort. Proteins were considered differentially expressed at |log_2_ fold change| > 0.5 and adjusted *P* < 0.05, unless otherwise specified.

#### Subtype-specific comparisons

For subtype-level analyses, we focused on tumor samples from the CPTAC dataset and applied separate modeling strategies for immune and cancer subtypes. Prior to modeling, the data were median-centered, and proteins with more than 20% missing values were excluded. For immune subtype comparisons, a linear model was constructed using both tumor purity (ESTIMATE_TumorPurity)^65^ and cohort membership as fixed covariates.

Differential expression was assessed using a leave-one-out (LOO) contrast approach. Modeling and empirical Bayes variance moderation were performed using Limma, and protein-level variance was further corrected using DEqMS via spectraCounteBayes based on PSM counts.

For cancer subtype comparisons, differential expression was performed within each cohort independently using tumor purity as the sole covariate; LOO contrasts were applied within each cohort to identify subtype-enriched proteins (|log_2_ fold change| > 0.5, adjusted *P* < 0.001).

### UPS protein–mutation quantitative trait locus (pQTL) analysis

UPS genes (n = 1,286) annotated in the Proteostasis Network (https://www.proteostasisconsortium.com/pn-annotation/) and represented in the CPTAC transcriptomic (n = 911) and proteomic (n = 971) data were analyzed. From a total of 19,932 gene mutations annotated across 10 CPTAC cohorts, 9,495 were observed in 11 or more tumor samples, the median across samples (**Supplementary Figure 4B**). For each UPS protein (Y) and each recurrent gene mutation (M), we fitted the linear model:

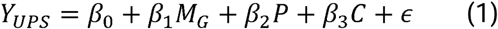

where P is tumor purity (ESTIMATE) and C is patient cohort. Wilcoxon rank-sum tests were also performed on UPS expression between mutated and wild-type groups across cohorts as a complementary analysis. Per-cancer associations were computed within individual cohorts. Raw p-values were adjusted using the Benjamini–Hochberg (BH) procedure, and the log_2_(mut/wt) fold change of the median or mean expression was calculated per data type.

### UPS stratification and survival analysis

Normalized protein abundance data for 922 UPS components, including 411 E3 ligases, were compiled from 11 CPTAC cancer types. Tumor samples were classified as UPS-high or UPS-low based on the upper and lower quartiles (75th and 25th percentiles, respectively) of normalized protein abundance within each cancer cohort, separately for each UPS component and tumor type. Clinical metadata, including overall survival (OS) time and vital status, were obtained from CPTAC sample annotations. Kaplan–Meier (KM) curves were generated using the *survival* and *survminer* R packages, and statistical significance was assessed using the log-rank test (*P* < 0.05). To evaluate independent prognostic value, multivariate Cox proportional-hazards models were fitted using the *coxph* function. Covariates included age, sex, and tumor purity^65^ (pan-cancer) or tumor purity alone (per-cancer). Hazard ratios (HR), 95% CIs, and *p*-values were reported, and adjusted *p*-values were computed using BH. Significant associations required log-rank *P* < 0.05, multivariate Cox *P* < 0.05, and |logHR| > 0.5. Analyses were performed in R (v4.2.2).

### Pathway analysis between high vs. low E3 protein groups

To investigate proteome-wide consequences of E3 ligase dysregulation, we performed differential protein expression analysis comparing tumors with high vs. low E3 protein abundance using the same quartile-based stratification (75th vs. 25th percentile). For each E3, log_2_-transformed protein abundance was modeled as:

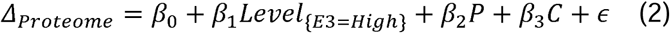

Linear models were fitted using DEqMS (v1.27.1)^63^ with empirical Bayes variance moderation. Differentially expressed proteins (adjusted *P* < 0.05, |log_2_FC| > 0.5) were submitted to gene set enrichment analysis (GSEA) using fgsea (v1.35.6)^66^ with pre-ranked lists based on the signed – log(*P*), against MSigDB gene sets via msigdbr (v25.1.1)^67^. Enriched pathways were defined at FDR < 0.05.

### DepMap gene co-dependency analysis

CRISPR-Cas9 gene-effect scores from DepMap Public 25Q2 (www.depmap.org) were filtered to CNS/Brain (n = 89) and Head and Neck (n = 77) cancer cell lines based on Oncotree lineage classifications. Pairwise Pearson correlations were computed across all genes using *Hmisc::rcorr* (v5.2.3)^68^; *p*-values were BH-corrected and signed enrichment scores defined as signed –log_10_(FDR). Pairs containing UBR5 or TRIM28 were extracted to build E3-centric co-dependency profiles. Ranked lists were used as input for pre-ranked GSEA (fgsea v1.35.6)^66^ against MSigDB^67^. Visualization used R (v4.5.1).

### UPS association with drug sensitivity in cell line models

Drug response data were obtained from PRISM Repurposing screens, which measure relative cell viability (AUC) across 578 cancer cell lines treated with 4,518 compounds^50^. We focused on cell lines annotated as Head and Neck or CNS/Brain lineages and matched them with normalized UPS protein abundance (or mRNA when protein was unavailable) from CCLE. For each UPS component, Spearman correlations were computed between expression levels and drug AUC scores within each lineage. Associations with *P* < 0.05 and |ρ| > 0.3 were considered significant.

## Ethics statement

This study did not involve the prospective recruitment of human participants, the collection of new human biospecimens, or any experiments in animals. All analyses were performed on de-identified proteogenomic datasets that had been previously generated, peer-reviewed, and released by established consortia and public resources, including CPTAC. Access to controlled CPTAC pan-cancer data was granted through dbGaP (phs001287.v16.p6) and the NCI Cancer Data Service. Because only de-identified secondary data were analyzed, no additional IRB approval was required at NYU Grossman School of Medicine.

## Acknowledgements

T.J.G.R. is an HHMI Gilliam Fellow (GT15758) and received funding from the NIH Institutional training grant in Cell Biology (T32GM136542). S.K. is supported by the NIH/NIGMS Pathway to Independence Award (K99GM155613) and the Weizmann Women’s Postdoctoral Career Development Award in Science. K.V.R. is thankful for funding from NIH/NCI (1U54CA263001-01A1). M.P. is a Howard Hughes Medical Institute Investigator. B.G.N. receives support from the Adelson Medical Research Foundation (AMRF). The authors thank the following for experimental feedback and critical discussion: Rodrigo Romero, Miguel Ramirez-Otero, Francisco J. Sánchez-Rivera, Rebecca Feltham, and Ngee Kiat (Jake) Chua.

## Author Contributions

T.J.G.R., K.V.R. and M.P. conceived and conceptualized the project. T.J.G.R. and K.V.R. developed the computational methodology. K.V.R. and M.P. supervised progress. T.J.G.R. and H.Z. analyzed protein expression profiles. T.J.G.R. and M.K. performed pQTL. T.J.G.R. completed survival, lineage-specific dependency and drug-sensitivity analyses and led the UBR5 and TRIM28 case studies. S.K., Y.K., G.R., D.F., B.G.N., and Y.M.S.F. provided intellectual input and advice on the study. T.J.G.R. wrote the original manuscript. T.J.G.R., K.V.R., and M.P. reviewed and edited the manuscript with input from all authors.

## Declaration of generative AI and AI-assisted technologies in the writing process

The authors used ChatGPT 5.o to improve writing clarity and syntax of this manuscript. The final version was thoroughly reviewed and approved by all authors.

## Supplementary Figure Legends

**Supplementary Figure 1.**
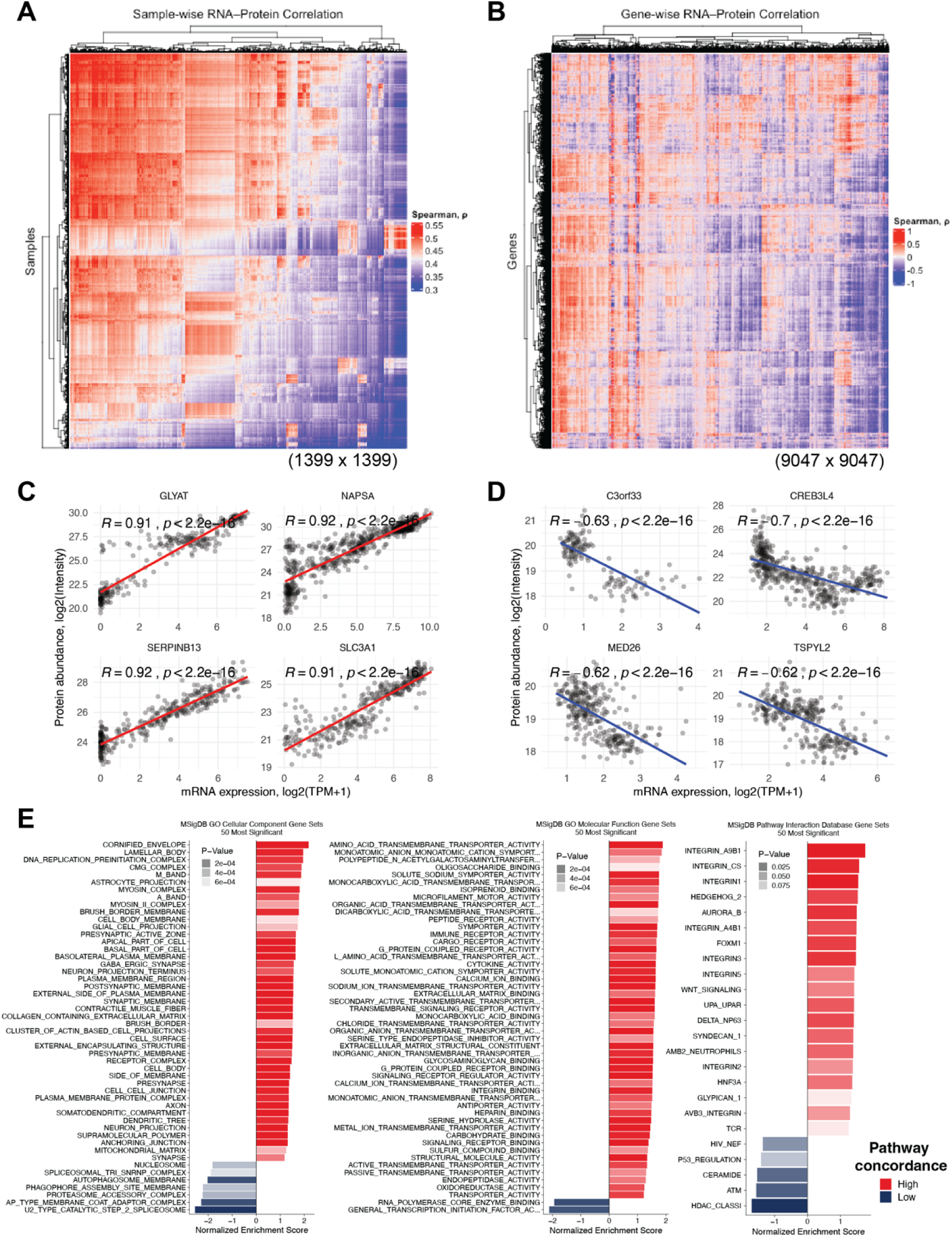
Sample- and gene-level mRNA-protein concordance in CPTAC. (A) Heatmap of sample-wise Spearman correlations between matched mRNA and protein profiles across CPTAC cohorts. (B) Heatmap of gene-wise Spearman correlations between mRNA and protein abundance. (C) Representative scatter plot of a gene with positive mRNA–protein correlation. (D) Representative scatter plot of a gene with negative mRNA–protein correlation. (E) Bar plots of pathway-level mRNA–protein concordance.

**Supplementary Figure 2.**
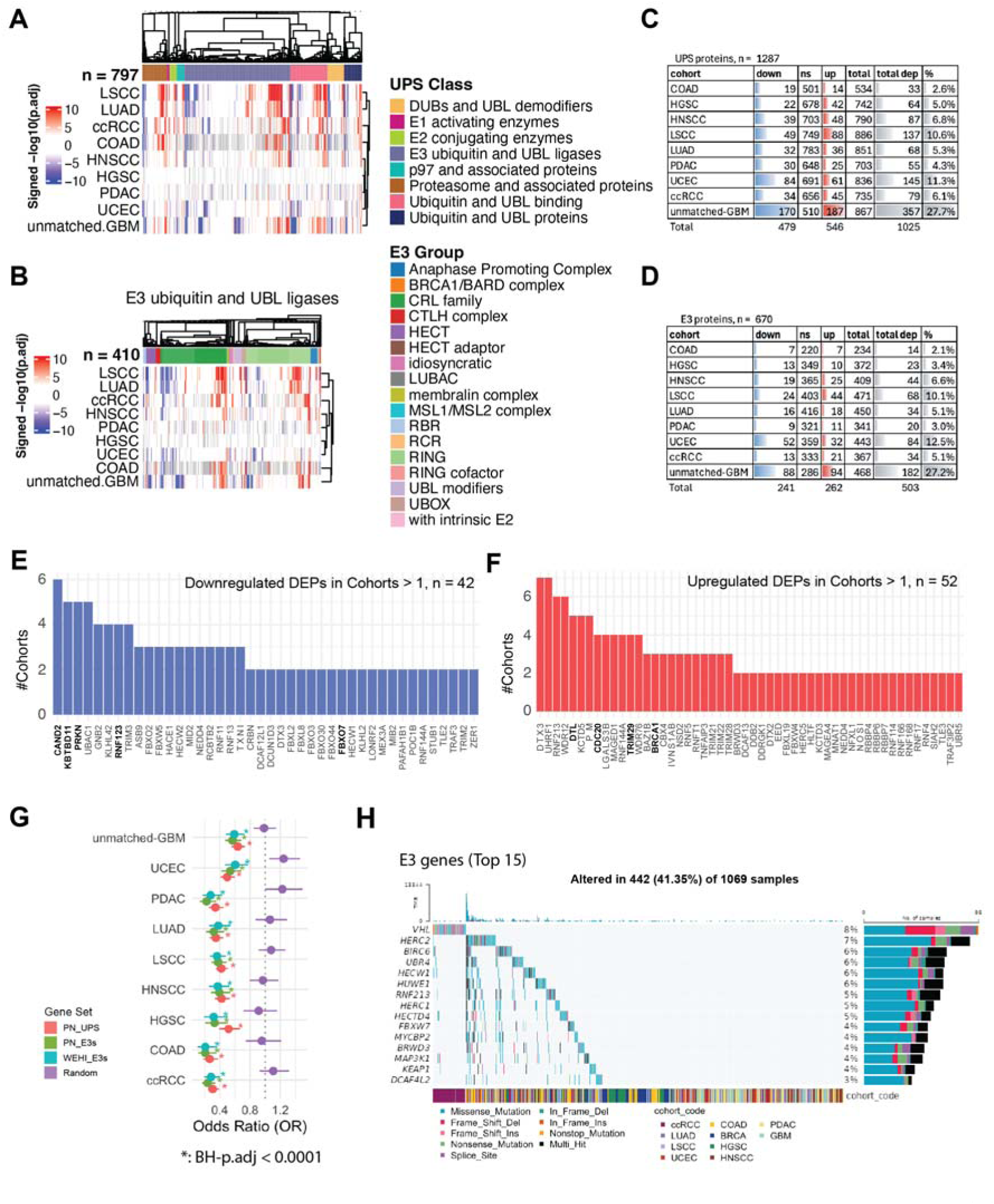
Differential expression landscape of UPS and E3 proteins. (A) Heatmap of signed –log_10_(adj. *P*) values for UPS-associated proteins (tumor vs. normal) across CPTAC cohorts. (B) Heatmap of signed –log_10_(adj. *P*) values for E3 ligases across cohorts. (C) Table summarizing the number/proportion of UPS-associated proteins significantly up- or downregulated in each cohort. (D) Same as (C) for E3 ligases. (E) Bar plot of significantly downregulated UPS proteins across cohorts. (F) Bar plot of significantly upregulated UPS proteins. (G) Fisher’s exact tests comparing dysregulation frequency among UPS-associated proteins versus randomly sampled protein sets. (H) Oncoplot of frequently mutated E3 ligases across CPTAC cohorts.

**Supplementary Figure 3.**
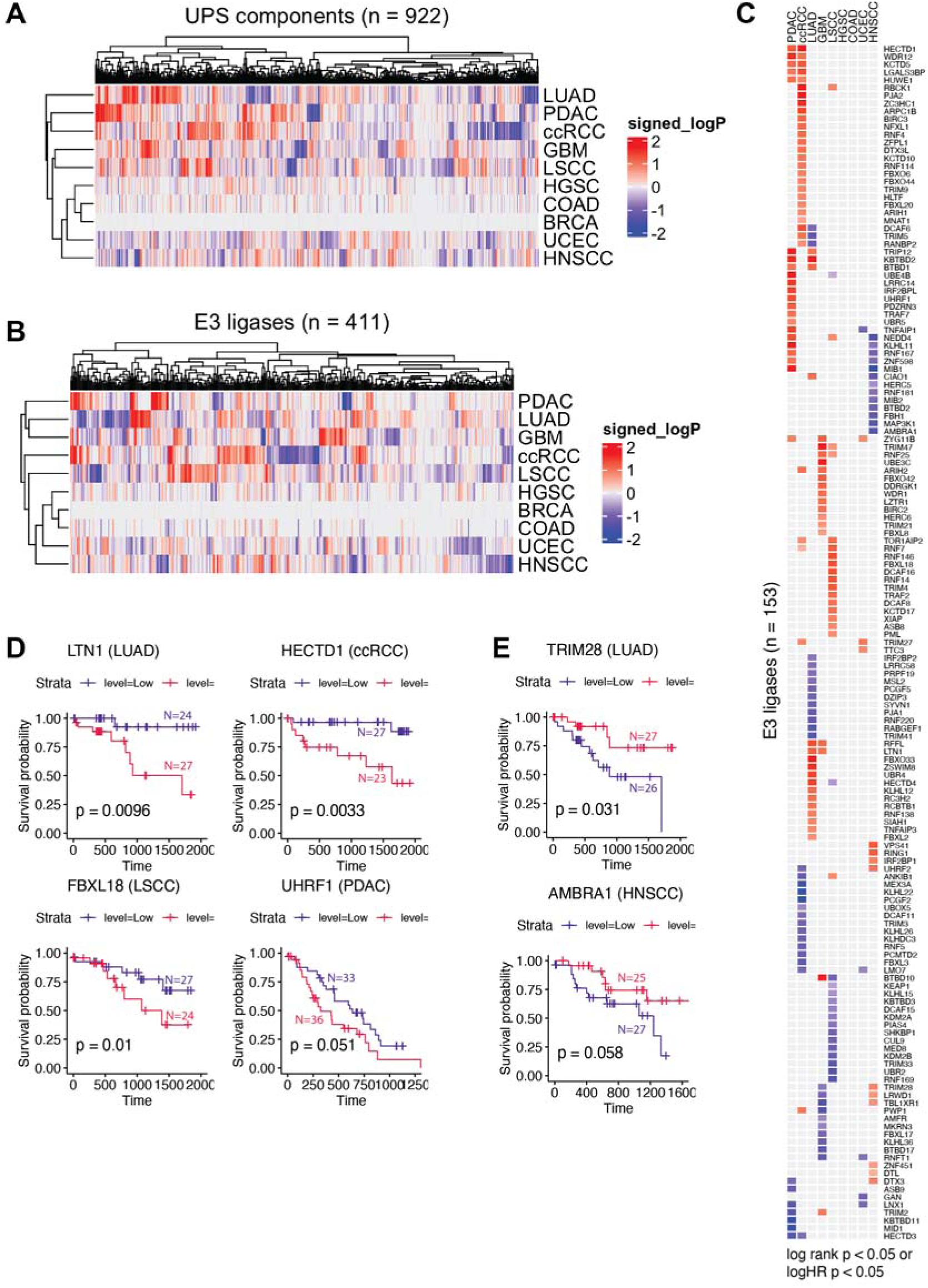
Survival associations of UPS components and E3 ligases. (A) Heatmap of pan-cancer overall-survival associations for UPS-associated proteins (Cox log-hazard ratio). (B) Same as (A) for E3 ligases. (C) Set of E3 ligases (n = 153) with significant overall-survival associations in at least one tumor type. (D) Representative Kaplan–Meier curve for an E3 with poorer survival when high. (E) Representative KM curves for E3s with improved survival when high.

**Supplementary Figure 4.**
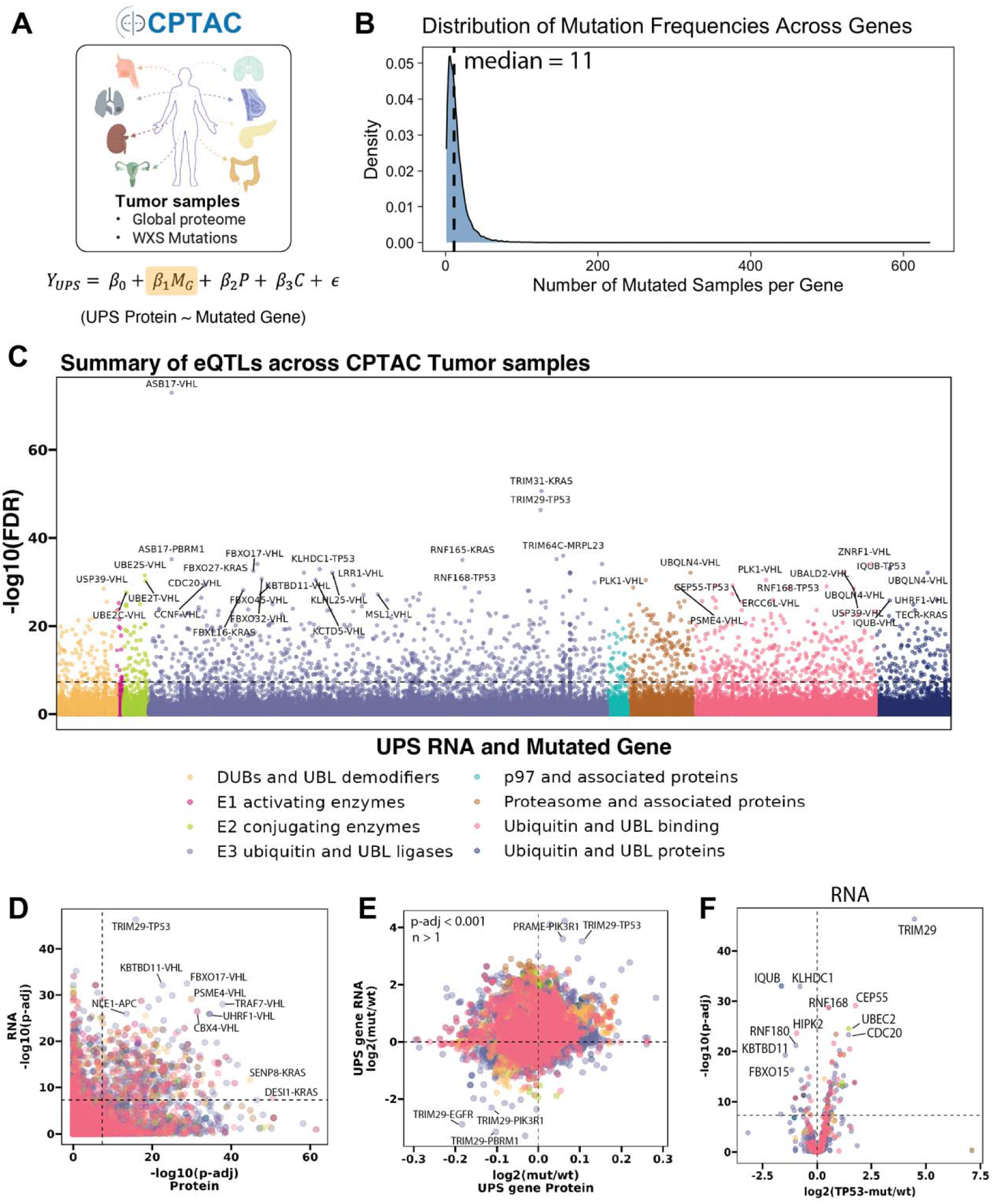
Expanded mutation-to-expression quantitative-trait-locus analysis. (A) Schematic of CPTAC datasets and the linear-modeling framework (Eq. 1). (B) Density plot of mutant-sample counts per recurrently mutated gene across CPTAC cohorts. (C) Manhattan plot of transcript-level mutation-to-expression QTL associations. (D) Scatter plot comparing –log_10_(adj. *P*) values from mRNA- vs. protein-level mutation associations for UPS genes. (E) Scatter plot comparing log_2_ fold changes from mRNA- vs. protein-level mutation associations. (F) Volcano plot of *TP53*-mutation effects on UPS-associated mRNA abundance.

**Supplementary Figure 5.**
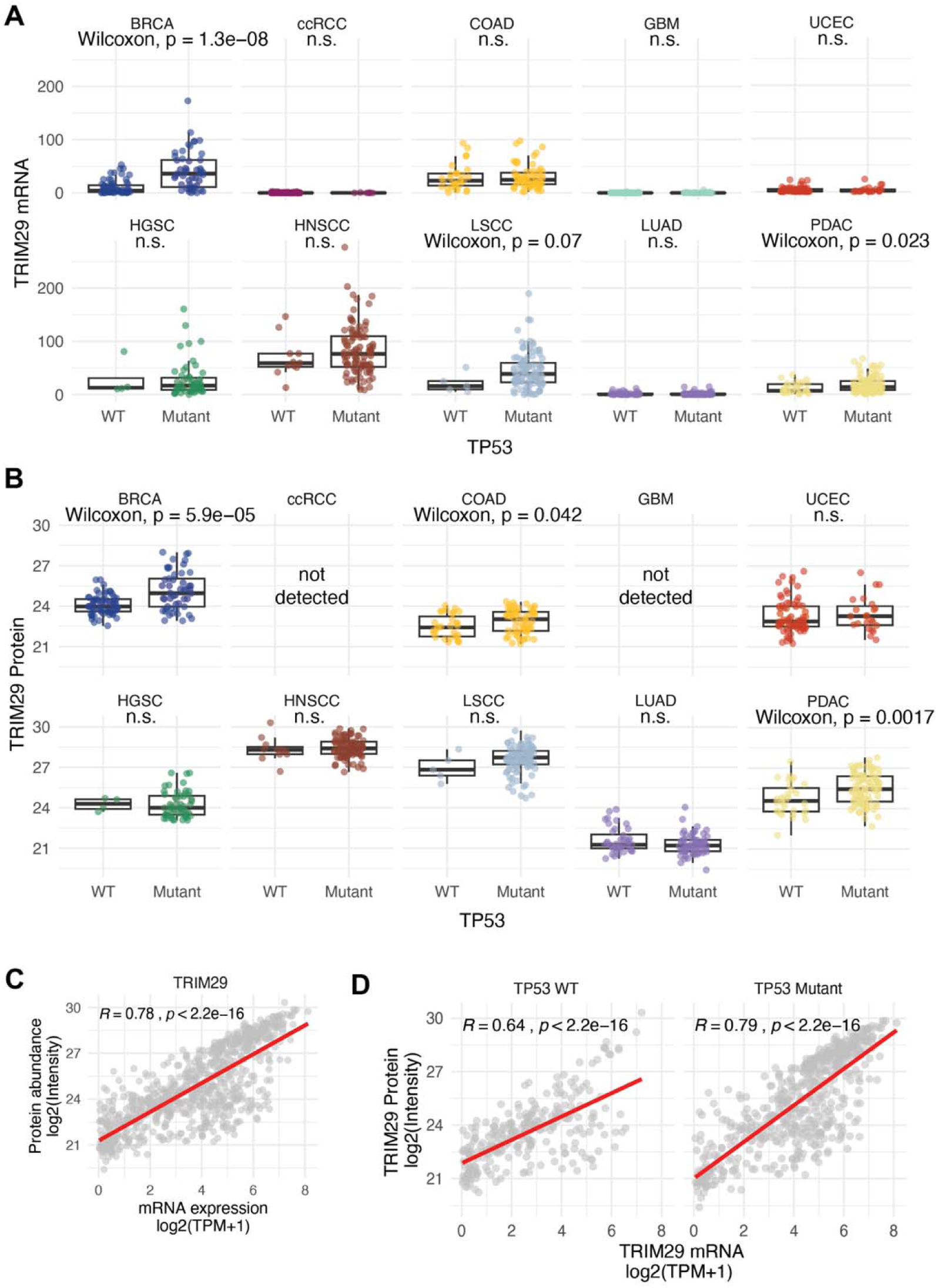
TRIM29 expression stratified by *TP53* mutation status. (A) Distribution of TRIM29 mRNA abundance in *TP53*-mutant vs wild-type CPTAC tumors. (B) Distribution of TRIM29 protein abundance, same stratification. (C) Scatter plot of TRIM29 mRNA vs protein abundance across CPTAC tumors. (D) Same as (C), stratified by *TP53* status.

**Supplementary Figure 6.**
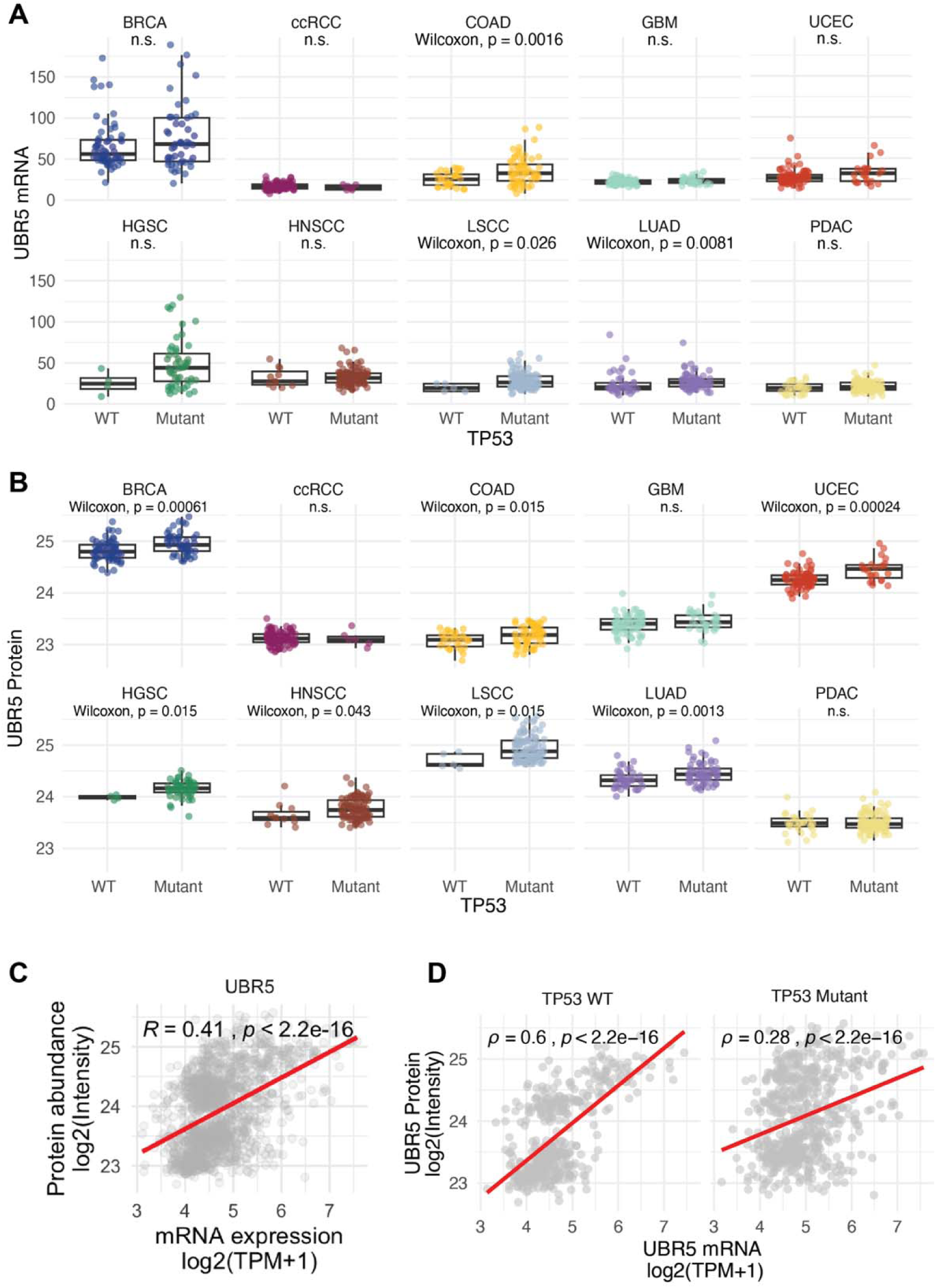
UBR5 expression stratified by *TP53* mutation status. (A) Distribution of UBR5 mRNA abundance in *TP53*-mutant vs wild-type CPTAC tumors. (B) Distribution of UBR5 protein abundance, same stratification. (C) Scatter plot of UBR5 mRNA vs protein abundance across CPTAC tumors. (D) Same as (C), stratified by *TP53* status.

**Supplementary Figure 7.**
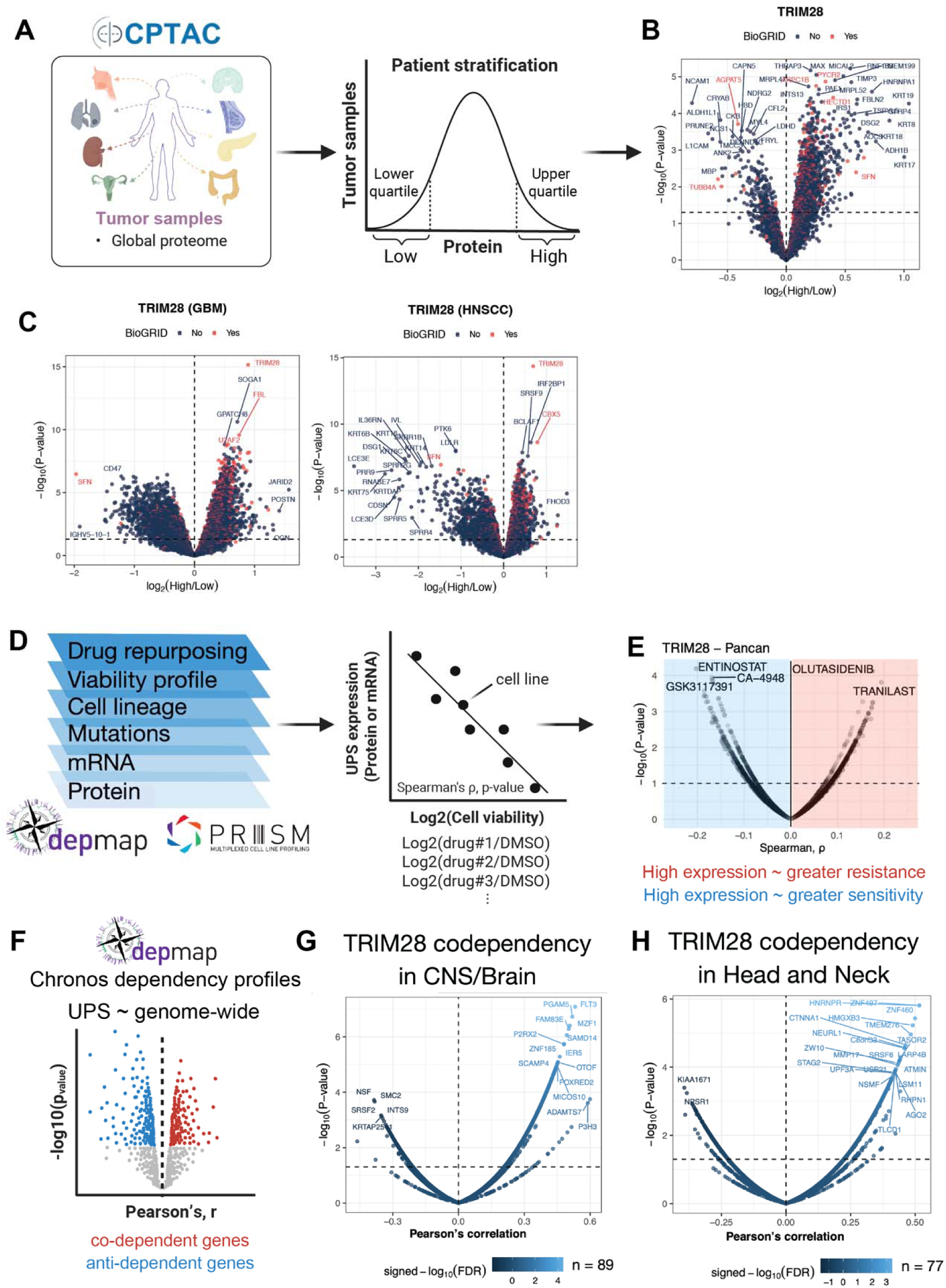
Differential expression, drug sensitivity, and dependency analyses for TRIM28. (A) Schematic of the differential analysis comparing TRIM28-high and TRIM28-low tumors across CPTAC cohorts. (B) Pan-cancer volcano plot of differential protein abundance (TRIM28-high vs low). (C) Lineage-specific volcano plots (GBM, HNSCC). (D) Schematic of the drug-sensitivity analysis. (E) Pan-cancer volcano plot of TRIM28–PRISM associations. (F) Schematic of the co-dependency analysis. (G) Volcano plot of TRIM28 co-dependency in CNS/brain. (H) Volcano plot of TRIM28 co-dependency in HNSCC.

**Supplementary Figure 8.**
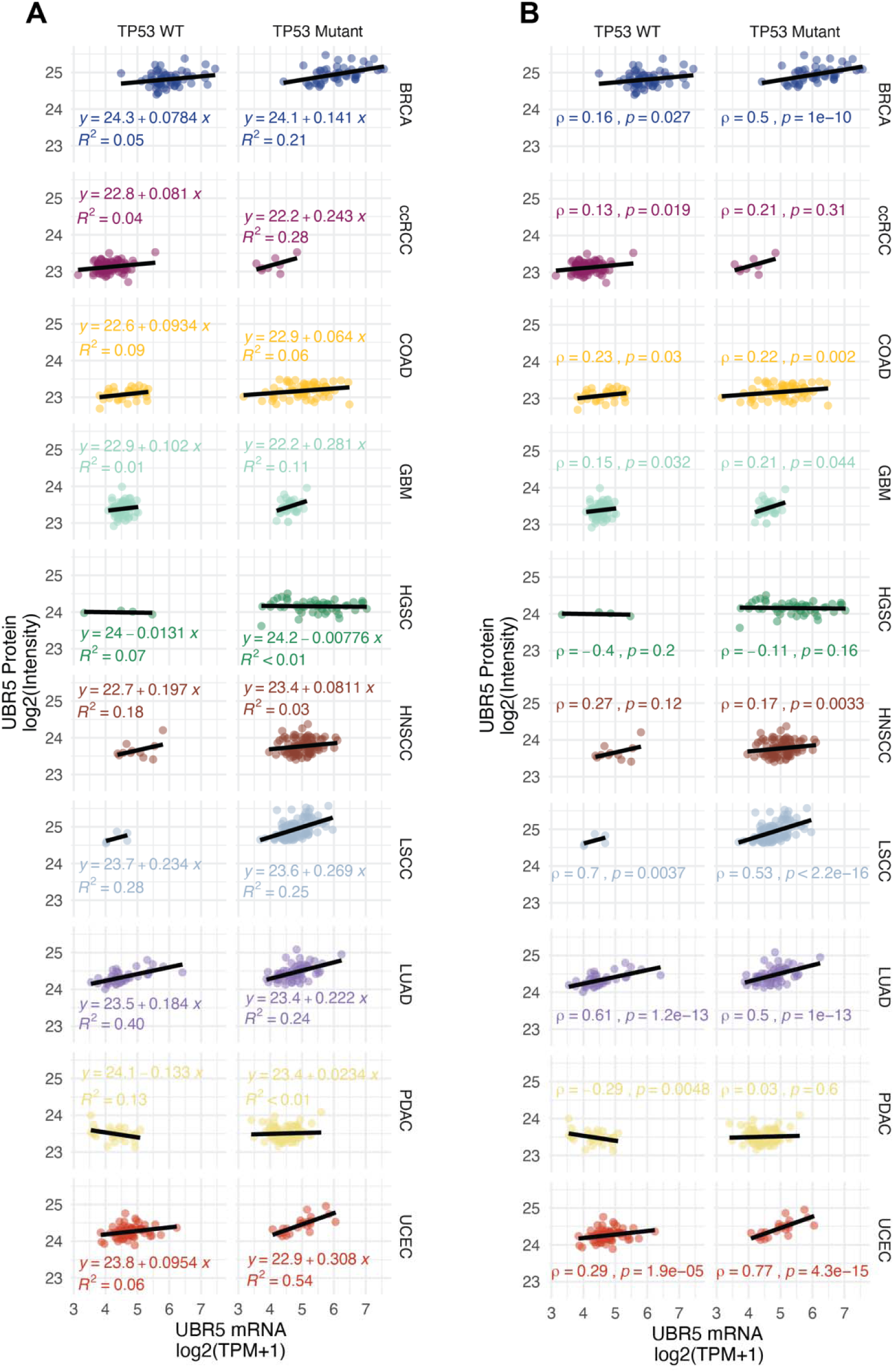
RNA–protein concordance for UBR5. (A) Linear regression of UBR5 mRNA vs protein abundance across CPTAC tumors. (B) Gene-wise Spearman correlation between UBR5 mRNA and protein abundance.

**Supplementary Figure 9.**
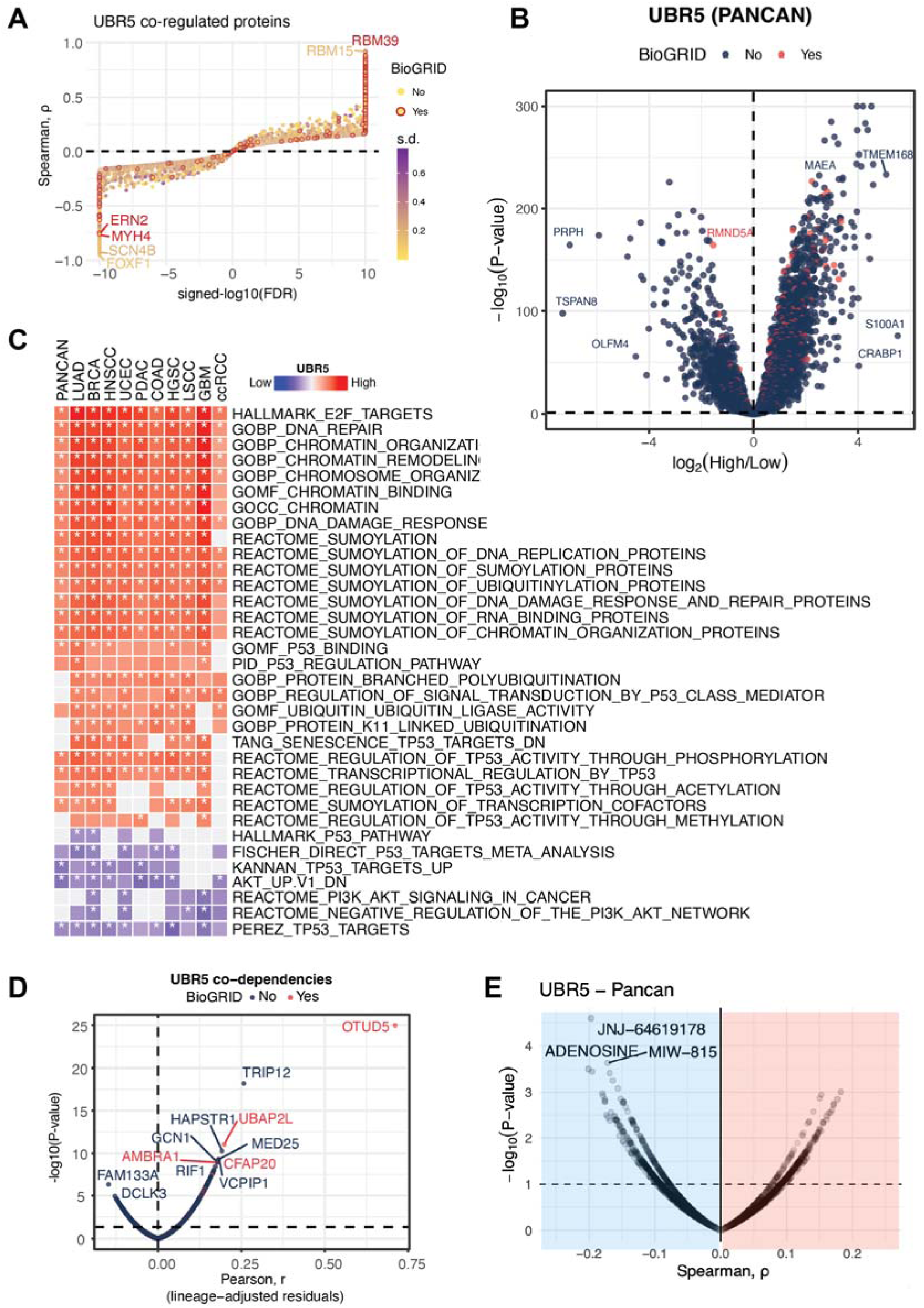
Pan-cancer UBR5 protein co-regulation and functional associations. (A) Pan-cancer UBR5 protein co-regulation profile (Spearman’s correlation). (B) Volcano plot of pan-cancer UBR5 protein co-regulation. (C) Heatmap of GSEA results from UBR5 protein co-regulation at pan-cancer and lineage-specific resolution. (D) UBR5 co-dependency profile from DepMap CRISPR Chronos with BioGRID interaction annotations. (E) Volcano plot of UBR5–PRISM drug sensitivity.

**Supplementary Figure 10.**
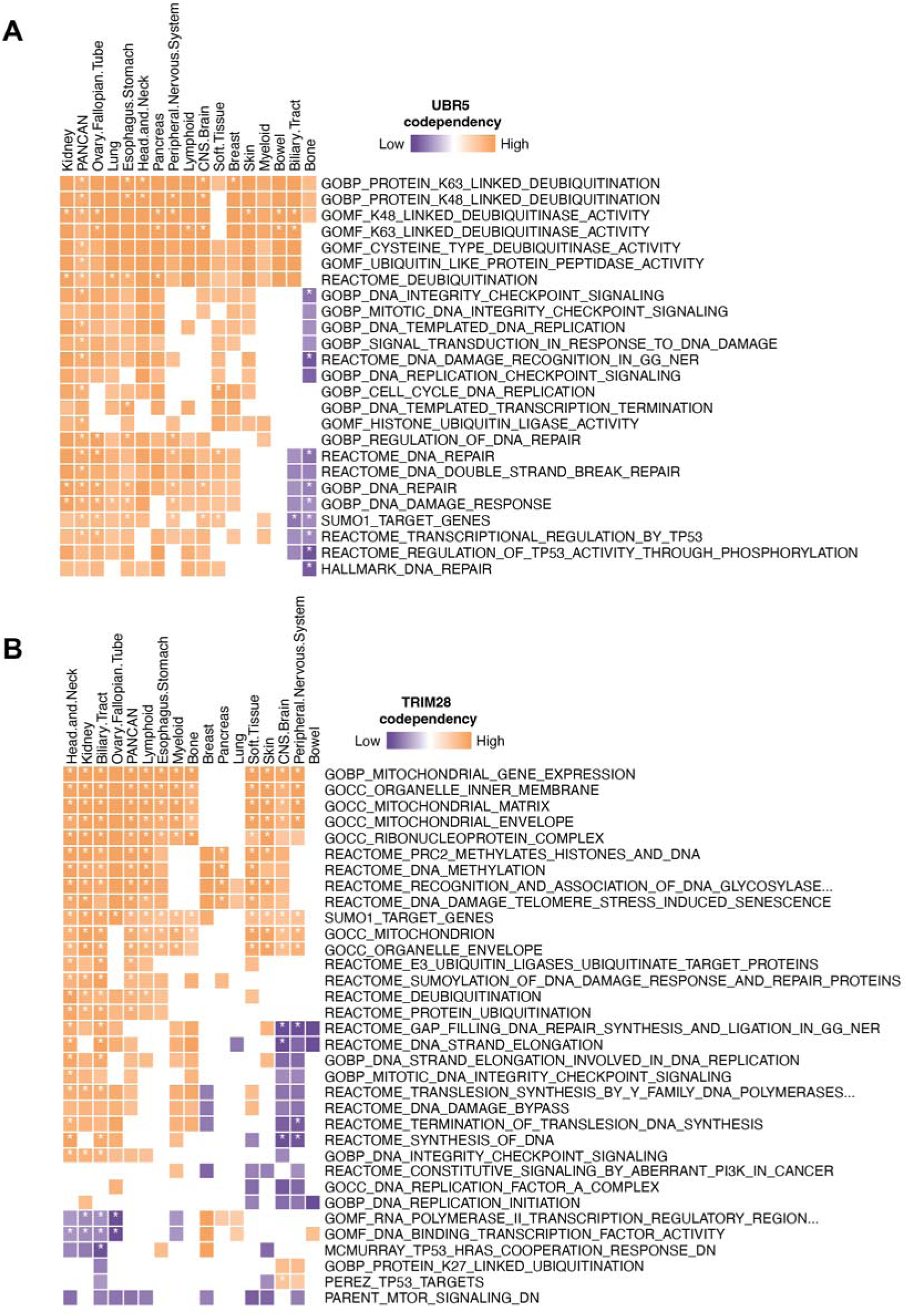
Pathway enrichment based on co-dependency profiles. (A) GSEA heatmap derived from UBR5 co-dependency profiles at pan-cancer and lineage-specific resolution. (B) GSEA heatmap derived from TRIM28 co-dependency profiles at pan-cancer and lineage-specific resolution.

## Supplementary Tables

**Supplementary Table 1**. Underlying data for specific main and supplementary figures, including mRNA-protein correlation and pQTL analysis.

**Supplementary Table 2.** Differential UPS protein expression across CPTAC cohorts (per-cohort tumor vs. normal contrasts; adj. *P*, |log_2_FC|).

**Supplementary Table 3.** UBR5 and TRIM28 co-regulation, GSEA, co-dependency, and PRISM drug-sensitivity results.

